# Prolonged tonic pain in healthy humans disrupts intrinsic brain networks implicated in pain modulation

**DOI:** 10.1101/740779

**Authors:** Timothy J. Meeker, Anne-Christine Schmid, Michael L. Keaser, Shariq A. Khan, Rao P. Gullapalli, Susan G. Dorsey, Joel D. Greenspan, David A. Seminowicz

## Abstract

Neural mechanisms of ongoing nociceptive processing in the human brain remain largely obscured by the dual challenge of accessing neural dynamics and safely applying sustained painful stimuli. Recently, pain-related neural processing has been measured using fMRI resting state functional connectivity (FC) in chronic pain patients. However, ongoing pain-related processing in normally pain-free humans remains incompletely understood. Therefore, differences between chronic pain patients and controls may be due to comorbidities with chronic pain. Decreased FC among regions of the descending pain modulation network (DPMN) are associated with presence and severity of chronic pain disorders. We aimed to determine if the presence of prolonged tonic pain would lead to disruption of the DPMN. High (10%) concentration topical capsaicin was combined with a warm thermode applied to the leg to create a flexible, prolonged tonic pain model to study the FC of brain networks in otherwise healthy, pain-free subjects in two separate cohorts (n=18; n=32). We contrasted seed-based FC during prolonged tonic pain with a pain-free passive task. In seed-based FC analysis prolonged tonic pain led to enhanced FC between the anterior middle cingulate cortex (aMCC) and the somatosensory leg representation. Additionally, FC was enhanced between the pregenual anterior cingulate cortex (pACC), right mediodorsal thalamus and the posterior parietal cortex bilaterally. Further, in the seed-driven PAG network, positive FC with the left DLPFC became negative FC during prolonged tonic pain. These data suggest that some altered DPMN FC findings in chronic pain could partially be explained by the presence of ongoing pain.

## Introduction

Functional connectivity (FC) has emerged over the past two decades as a technique to investigate the functional anatomy of human brain networks and the effects of psychological, pathological and perceptual manipulations, therapeutic treatments, and disease states on brain function [5;7;21;42;78]. The FC of the descending pain modulatory network (DPMN) has been well-studied during simultaneous experience of phasic painful heat stimuli, concurrent painful heat and distracting stimuli, during mind-wandering, and in the context of placebo analgesia [6;28;46;52;71;96]. The DPMN frequently includes the subgenual and pregenual anterior cingulate cortices (sACC and pACC), anterior midcingulate cortex (aMCC), and periaqueductal gray (PAG), whether studied at rest, in a neutral environment, or during experience of painful stimuli [28;45;51;52;71;99;100]. Demonstrating the potential clinical importance of the DPMN, enhanced FC within the DPMN during placebo analgesia and motor cortex stimulation is positively related to magnitude of analgesia experienced [6;28;33;64]; additionally, there is a reduction of FC of the DPMN in chronic pain patients compared to pain-free controls [52;108]. In healthy individuals, coactivation of the PAG and pACC during the experience of phasic pain coupled with analgesic cognitive manipulations, such as placebo, results in enhanced FC between these regions accompanied by a reduction in perceived pain intensity [6;28;46;96].

While several studies have evaluated the network’s FC during phasic painful stimuli there is a scarcity of studies exploring FC of the DPMN during prolonged tonic pain [6;46;86;96]. Persistent pain states, such as those encountered in chronic pain syndromes, display unique perceptual dynamics and modeling prolonged tonic pain in healthy subjects is a critical intermediate step in understanding the neurophysiology of chronic pain disorders [4;31].

To probe the connections of the DPMN during a prolonged tonic painful stimulus in a preclinical human pain model, we acquired resting state fMRI scans in pain-free subjects before and after exposing them to a potent topical capsaicin-heat pain (C-HP) model [2;64], thereby capturing pain-free and prolonged tonic pain resting states within a single imaging session.

In this report, we focus our analysis on the FC of the nodes of the descending pain modulatory network: aMCC, sACC, pACC, and PAG [6;28;33;71;93]. We predicted that prolonged tonic pain would disrupt the coupling between the pACC and PAG since disruption in FC between pACC and PAG occurs in chronic pain disorders and the pACC displays reductions in local FC in chronic pain [41;42;49;53;102;105]. Additionally, we predicted enhanced FC of pACC to aMCC and diminished FC between PAG and sACC during prolonged tonic pain. Evidence supporting these predictions would support the role of prolonged tonic pain as a factor in modulations of resting state FC in people with chronic pain.

## Methods

### Overview

We report results from two studies conducted at University of Maryland Baltimore (UMB) from October 2011 until December 2015. Study 1 set out to establish the effects of a prolonged tonic pain stimulus lasting several minutes on the functional organization of the human brain. In the first experiment, we conducted experiments provoking pain in 18 healthy subjects (10 M; 2 left-handed) aged 23 to 61 (median = 30.5) by applying 10% capsaicin cream under a warm thermode on their left leg, which we term the capsaicin-heat pain (C-HP) model to distinguish it from previous models using lower concentration capsaicin creams [2; 13; 64; 70]. We conducted a screening session to eliminate subjects who did not develop sufficient heat allodynia during the C-HP [54]. Eligible subjects then took part in an MRI session which was separated by ≥13 days (median=39.5 (range=13 to 88)) from the screening session. Using the C-HP model we maintained a mild to moderate pain intensity with a 39°C (n = 11), 40 °C (6) or 41 °C (1) thermode. The temperature selected for each participant was based on that individual’s thermal heat pain sensitivity evaluated at screening just prior to the C-HP exposure.

In study 2, 40 subjects (17 M; median age: 24; range 20-39) underwent an MRI in which we employed the C-HP model with a 38°C (n = 4), 39 °C (2), 40 °C (6), 41 °C (8), or 42 °C (20) thermode. From this group of subjects, 3 males were excluded from the current report because they did not report pain ratings greater than an average rating of at least 10 out of 100 during the last 2 minutes of exposure to the C-HP model (C-HP temperatures for these subjects were all 42 °C). A further 4 males and 1 female were excluded due to having greater than 0.5 mm motion framewise displacement in at least 10% of the functional MR time series (1 at 40 °C, 1 at 41 °C, 2 at 42 °C). Resting state fMRI results from study 1 and 2 are pooled together and use the same MRI sequence and protocol, excepting that scans were acquired at different resolutions (1.8 x 1.8 x 4 mm^3^ versus 3 mm^3^ isotropic) and different head coils (12- versus 32- channel). All subjects provided written informed consent, and all procedures were approved by the UMB Institutional Review Board for the Protection of Human Subjects.

### Eligibility Criteria

In study 1 exclusion criteria were: pregnancy; history of brain injury with any period of unconsciousness; illicit, or prescription opioid, drug use; current pain or history of chronic pain; history of cardiac, renal, hepatic, or pulmonary function disorders; history of cancer; ambidextrous [67]; hospitalized for a psychiatric disorder within last 12 months; pain intensity rating less than 21 on a 0-100 NRS while exposed to the C-HP model. Illicit drug use was determined with a urine drug screen for marijuana, cocaine methamphetamine, amphetamines, ecstasy, heroin, phencyclidine, benzodiazepines, methadone, barbiturates, tricyclic antidepressants or oxycodone (First Check™).

In study 2, in addition to eligibility criteria for study 1, we excluded left-handed subjects, any individual with any diagnosis of psychological or neurological disorder or subjects taking any psychoactive medications (by self-report). However, in study 2 we did not exclude any individual based on sensitivity to the C-HP model.

### Psychophysics and Psychological Questionnaires

During the initial session of each study, we measured subjects’ warmth detection thresholds (WDTs) and heat pain thresholds (HPTs) with a Medoc stimulator (Pathway; Medoc; Ramat Yishai, Israel) using the method of limits [35]. We placed the 3×3 cm contact area stimulator on the lower left foreleg at a baseline temperature of 32 °C. A program increased the temperature at a ramp of 0.5 °C/s until the subject pressed a mouse button. We instructed the subject to press the button when they “felt a change in temperature” for WDTs or when the warmth “becomes painful” for HPTs. At a single site, we measured four trials for WDTs and HPTs. We took the average of the last three threshold determinations for each subject. In study 2, the protocol for HPTs started with a baseline of 30 °C, to accommodate sensitivity changes after capsaicin exposure.

### Capsaicin-Heat Pain Model

In order to produce a safe, sustained painful experience, we treated subject’s lower left foreleg with one gram of 10% capsaicin cream under a Tegaderm™ bandage [2; 64]. To control the exposure area, we applied the cream within a 2.5 cm^2^ square cut into a Tegaderm™ bandage. After 12 minutes of exposure – long enough for the capsaicin cream to reach saturating concentrations at the intraepidermal nerve fiber endings – we placed the thermode over the topmost bandage at the designated temperature [34]. During study 2 the incubation period was increased to 15 minutes and the thermode was placed on the subject’s leg during incubation, held at 32 °C. Target temperatures used were tailored for each subject between the subject’s WDT and HPT determined prior to capsaicin exposure. Subjects rated pain intensity on a numerical rating scale (NRS) with verbal anchors on one side, and numbers ranging from 0 to 100 in increments of 10 [36]. In study 1, subjects provided pain intensity ratings every 30 seconds for 10 minutes after application of the thermode. Subjects reporting average NRS pain 30 out of 100, and tolerating the C-HP model, were eligible for MRI sessions. During study 2 subjects provided pain intensity ratings every minute for 35 minutes during the entire capsaicin exposure. At the end of the exposure period, we removed the bandages and capsaicin with an isopropanol swab. This C-HP procedure does not cause tissue damage [66].

### fMRI recording

We recorded fMRI in a 3-T Tim Trio scanner (Siemens Medical Solutions, Malvern, PA) using a 12-channel (study 1) or 32-channel (study 2) head coil with parallel imaging capability. For resting state scans during study 1, we used a gradient echo single-shot echo-planar-imaging (EPI) sequence with 30 ms echo time (TE), 90° flip angle and 2500 ms repetition time (TR) providing T2*-weighted volumes in 36 interleaved, 4 mm slices (no gap) with an in plane resolution of 1.8 mm^2^. For study 2, we used identical parameters except that we collected 44 interleaved, 3 mm slices (no gap) with an in-plane resolution of 3.0 mm^2^. During both resting state scans subjects fixated on a crosshair for 8 min 12.5 s providing 194 functional volumes. For anatomical reference, we acquired a 3-dimensional T1 magnetization-prepared rapid gradient echo (MPRAGE) volumetric scan with 2.9 ms TE, 2300 ms TR, 900 ms inversion time (TI), flip angle 9°, 144 slices, axial slice thickness 1.0 mm and 0.9 mm^2^ in-plane resolution over a 23-cm field of view for 13 of 14 subjects. We used a different MPRAGE protocol for the last subject of study 1 and all subjects of study 2: 2.9 ms TE, 2300 ms TR, 900 ms TI, flip angle 9°, 176 slices, sagittal slice thickness 1.0 mm and 1.0 mm^2^ in-plane resolution over a 25.6-cm field of view. Since structural scans were used for anatomical reference and display only, this difference in anatomical acquisition did not influence the results.

### fMRI Session Protocol

During study 1 and 2 MRI sessions, we evaluated the subjects’ WDTs and HPTs in the MRI environment. During the resting state scan subjects were told: “Please stare at the plus sign, do not move and do not fall asleep. You may let your mind wander.” Then, we increased the thermode temperature to the predetermined target, while the subject fixated on the cross hair for the duration of the scan. After the scan, subjects remained in the scanner with the thermode in place and rated their pain intensity every 30 seconds for two minutes on a 0-100 NRS.

### Statistical Methods

Effects of time on sensory detection thresholds (WDTs and HPTs), after exposure to the C-HP model, were evaluated using a linear mixed model with time as a fixed factor and subject as a random factor. In different models, either baseline HPT or baseline WDT were using as a control to compare to HPTs taken at 25, 50 and 75 minutes after capsaicin removal. Multiple comparisons were corrected using Tukey’s HSD. The R package ‘anova’ was used to derive F-stats for the overall model. We conducted all statistical tests using R 3.4.1.

### Resting state fMRI data analysis

All preprocessing of resting state fMRI scans used the afni_proc.py python script for Analysis for Functional NeuroImaging (AFNI). The first three volumes were automatically removed from the functional scan series by the MRI scanner to allow for signal equilibration. We used 3dToutcount to determine volumes where more than 10% of the time points are outliers. Outliers were defined in relation to the median absolute deviation of the signal time course (see afni.nimh.nih.gov/pub/dist/doc/program_help/3dToutcount.html for outlier definition). Each TR was slice-time corrected and aligned to the top slice, collected midway through the TR. Then, each functional time series was detrended and spikes quashed with 3dDespike. Before aligning the anatomical scan to the functional scan, the skull was removed from each individual anatomy using 3dSkullStrip. We used 3dAllineate via the align_epi_anat.py python script to align the anatomy to the third functional volume acquired. After alignment, the anatomical volume was warped to Talairach atlas space and normalized to the ICBM452 brain using @auto_tlrc. Next, 3dAllineate applied the 12-parameter affine warping matrix, determined during normalization of the anatomical to Talairach space, to the registered functional volumes. Each 4D dataset was bandpass filtered between 0.008 and 0.1 Hz. We created individual subject functional masks from the registered normalized EPIs. Estimated individual maps of white matter (WM), cerebral spinal fluid (CSF) and gray matter (GM) were created for each subject using 3dSeg. We applied spatial blurring using the iterative program 3dblurtoFWHM with a filter of full width half maximal of 6 mm. This level of spatial smoothing was consistent with the group average estimate of noise smoothness in the functional time courses as estimated by 3dFWHMx (Study 1: session 1: x = 5.9 mm y = 5.9 mm z = 6.0 mm. Study 2: x = 6.1 mm y = 6.1 mm z = 6.0 mm).

In the subject-level regression model regressors of no interest included a binary regressor of outlier volumes and volumes with motion exceeding 0.5 mm, signal derived from the eroded WM and CSF mask, demeaned motion parameters (motion in x, y and z planes and rotation about the x, y and z axes), and their first order derivatives. For each subject the regressor of interest was the seed time course for that particular seed and session. The seed regions of interest were 5 mm radius spheres centered within the bilateral anterior midcingulate cortex (aMCC: +/- 6, 32, 22), pregenual anterior cingulate cortex (pACC: +/-6, 40, 6), subgenual anterior cingulate cortex (sACC: +/-6, 28, -4); and a subject-specific anatomically drawn seed for the periaqueductal gray (PAG; center of mass: 0, 29, -7) (Supplemental Figure 1). The PAG seed was drawn to encompass all gray matter surrounding the cerebral aqueduct.

### Group Level fMRI Analysis

For group level analysis, we used AFNI’s linear mixed-effects modeling program 3dLME [15]. A two-factor model focused on the change in state between the control, pain-free and the prolonged tonic pain resting states with the C-HP model inducing heat allodynia (factor levels: control and pain) to arrive at pain-control contrast maps and mean seed-driven intrinsic connectivity network (ICN) maps. For pain intensity covariation with seed-driven functional connectivity (FC), we used AFNI’s 3dttest++. For brain-wide non-significant results, we present the corresponding minimum false discovery rate adjusted q-value [91]. We implemented a minimal voxel-wise p-value threshold of 0.005 for covariate and contrast analyses as well as mean ICN maps derived from the PAG seed and 0.001 for mean ICN maps derived from bilateral pACC, sACC and aMCC seeds.

For voxel tables detailing contrasts and covariate results, we implemented an initial voxel-wise threshold of 0.005, while we thresholded all other voxel tables at an initial voxel-wise threshold of 0.001. In order to elucidate the coordinates of local maxima within large clusters, we iteratively reevaluated statistical maps after reducing the p-value threshold of each map by a factor of 10 (e.g. 0.001 to 0.0001). We chose a p-value threshold of 0.005 since this corresponds to a minimum false discovery proportion of 0.067 [19]. To correct for multiple comparisons, we estimated the spatial autocorrelation function of the residual noise of the BOLD signal within our analysis mask using 3dFWHMx and used the resulting parameters with 3dClustSim to calculate cluster extent criteria (CEC) for both 3dLME and 3dttest++ resultant maps. We further restricted the analysis to four anatomically-based masks encompassing cortical gray matter, cerebellum, subcortical gray matter, and brainstem (Supplemental Figure 2). We calculated CEC for p-values of 0.005 and 0.001 for each anatomical compartment separately since each compartment was assumed to behave differently (e.g. the thalamus and brainstem are assumed to have smaller foci or activation than the cerebral cortex) (Supplemental Table 1).

## Results

### Psychophysical and Perceptual Response to Capsaicin-Heat Pain (C-HP) Model During Screening Session

All subjects included in this analysis reported hot-burning pain in response to the C-HP model. For subjects in study 2 (n=32), the time course for pain intensity ratings during the screening session shows an ever-increasing trend during the C-HP exposure, which responded positively to 0.5° C step increases at each arrow (Fig. 1A). The thermode temperature for the MRI scan was tailored for each subject and, once increased to the target temperature, remained stable during the MRI sessions. Further, in study 2 (n=32), exposure to the C-HP model induced profound heat allodynia which lasted at least 75 minutes after C-HP removal (F=11.28; p<0.001) (Fig 1B). HPTs after capsaicin removal were all significantly lower than baseline HPTs (t≤ -7.8; p≤10^-8^) and the first HPTs after C-HP removal were lower than baseline WDTs (t=-4.5; p=8.4×10^-5^), demonstrating profound hypersensitivity to heat after C-HP exposure. Using SFMPQ-2 descriptors, most subjects described the pain as throbbing, aching, heavy, tender, shooting, stabbing, sharp, piercing, sensitive to touch, hot-burning and tingling (Fig 1C). After removing the descriptor ‘hot-burning’ from the analysis, we found a significant effect of pain descriptor class (F=9.17; p<0.0001) and no effect of sex on subjective ratings on the SFMPQ-2 (F=0.18; p=0.67). Subjects rated both intermittent and continuous pain descriptors higher than either neuropathic or affective pain descriptors (Continuous>Affective: t=4.0; p<0.001; Intermittent>Affective: t=4.0; p<0.001; Continuous>Neuropathic: t=4.1; p<0.001; Intermittent>Neuropathic: t=4.1; p<0.001). There was no difference in ratings between continuous and intermittent descriptors and neuropathic and affective descriptors (t<0.13; p>0.98).

**Figure 1.**
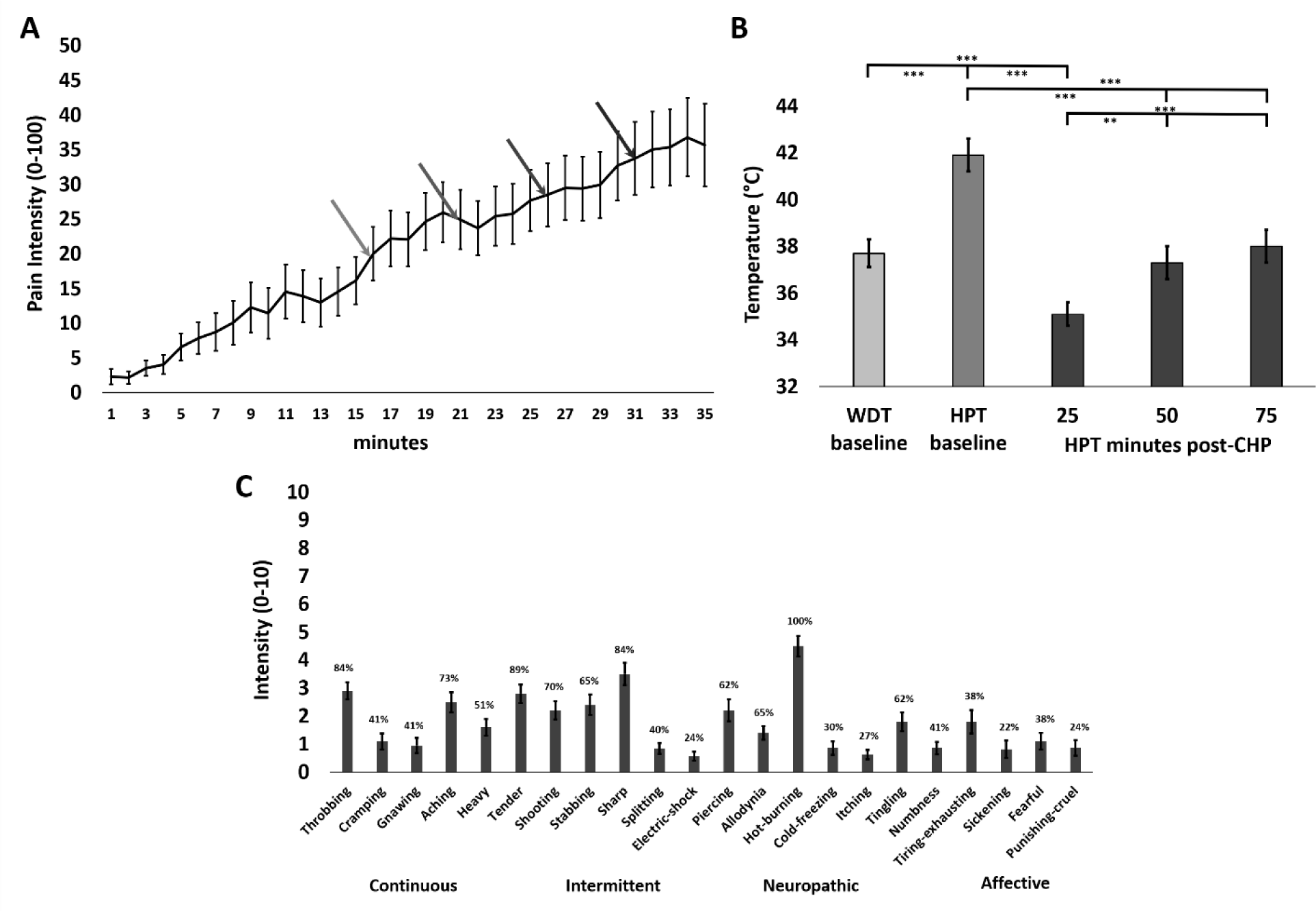
**A)** Ratings of prolonged tonic pain intensity during C-HP, the first arrow is when the thermode increased to the target temperature and the temperature increased 0.5°C at each subsequent arrow (n=32). **B)** Warmth detection thresholds (WDT) and heat pain thresholds (HPT) taken before and after exposure to the C-HP model (n=32); ** - p = 0.0002; *** - p < 0.0001. **C)** Mean intensity of pain descriptors from the SFMPQ-2, where percentages above each bar represent percentage of subjects endorsing pain descriptor with any non-zero rating. All error bars are SEM.

### Resting State Functional Connectivity: Topography during pain-free and prolonged tonic pain resting states

We used seed-driven methods to investigate the effect of prolonged tonic pain on the ICN of the DPM network, a group of structures proposed to be important in pain modulation. The DPMN includes the ACC and the PAG [6; 17; 28; 71; 93]. To investigate this network, we placed seeds in the PAG, sACC, pACC and aMCC. We interrogated the ICNs during a control state with subjects passively focusing on a crosshair and a prolonged tonic pain resting state with subjects experiencing the C-HP model on their left leg while passively focusing on a crosshair.

The pain-free ICN map derived from the bilateral aMCC displayed several large clusters of correlated signal (positive FC) across the brain. The first cluster consisted of a very large region of positive FC spreading from the bilateral aMCC into the lateral frontal poles (Figure 2A and Supplemental Table 2: Cluster 1). The second cluster was focused in the bilateral caudate and spread throughout the bilateral basal ganglia and thalamus as well as into the bilateral dorsal anterior and posterior insula (Cluster 2). A large cluster of anticorrelated signal was found in the left cerebellum as well as into the left fusiform and middle temporal gyrus (MTG) (Cluster 3). Furthermore, clusters of anticorrelated signal (negative FC) were found in the bilateral superior parietal lobules (SPLs) (Clusters 5 and 7). A significant cluster of negative FC was found in the rostral medulla (Cluster 19).

**Figure 2.**
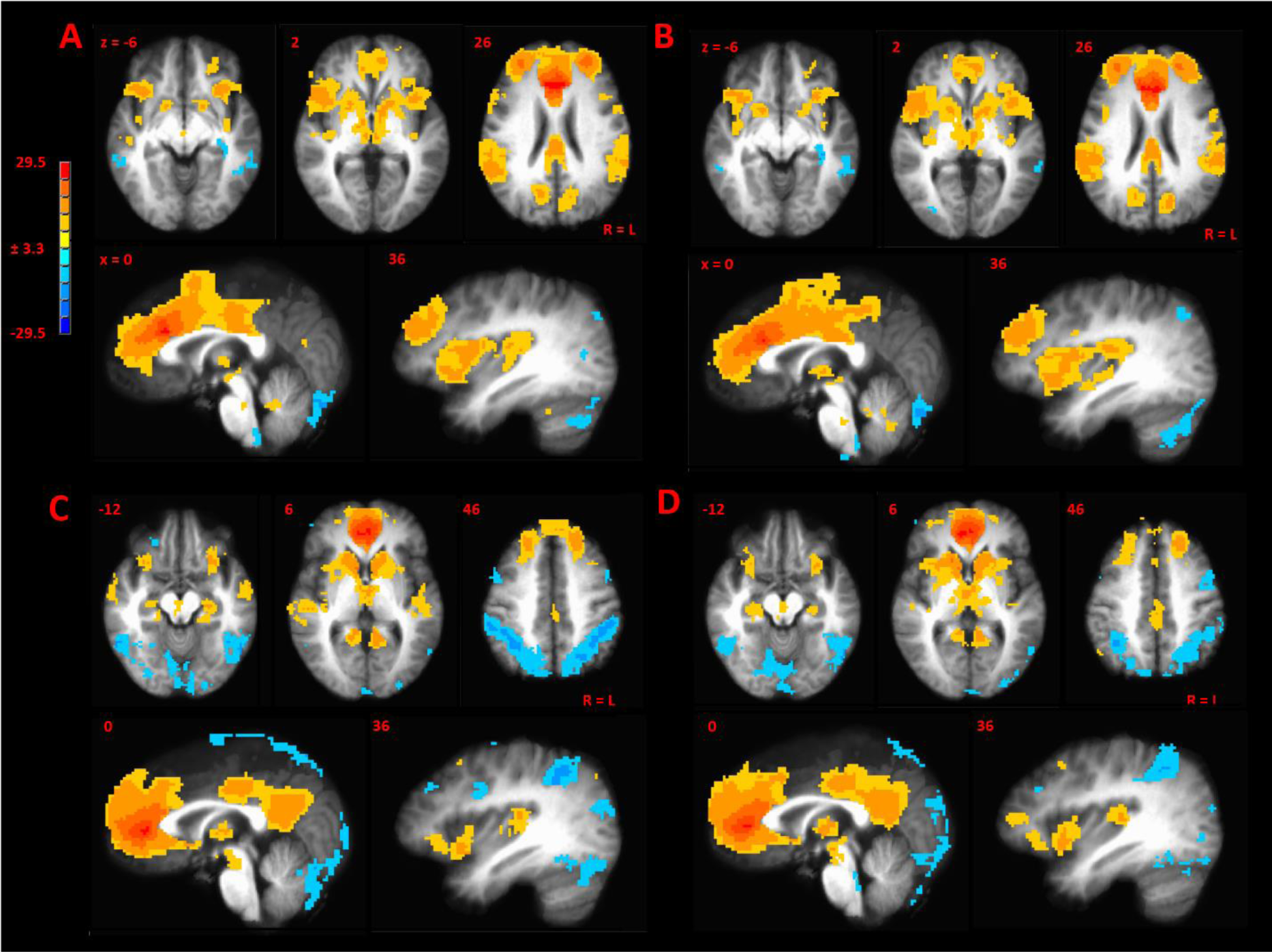
**A)** Mean seed-driven functional connectivity (SDFC) map from the aMCC seed during the control state. **B)** Mean SDFC map from the aMCC seed during prolonged tonic pain. **C)** Mean SDFC map from the pACC seed during the control state. **D)** Mean SDFC map from the pACC seed during prolonged tonic pain. R=L; Bar is in Z-units.

During prolonged tonic pain, aMCC FC was similar to the ICN map derived during the control state. A large cluster of positive FC surrounding the seed was present throughout the aMCC projecting in the bilateral frontal gyri (Figure 2B and Supplemental Table 3: Cluster 1). This cluster of positive FC spread bilaterally into the caudate and thalamus extending significant peaks in the bilateral middle and posterior insula as well as into the inferior parietal lobules (IPLs) (Cluster 1). A large cluster of negative FC spread throughout the cerebellum bilaterally as well as bilateral occipital gyri (Cluster 2). Additional clusters of negative FC included the left SPL (Cluster 4) and left parahippocampal gyrus (PHG) (Cluster 5). Again, a significant cluster of negative FC was found in the rostral medulla (Cluster 11).

In the control ICN map positive FC derived from the bilateral pACC spread bilaterally into the caudate nuclei, dorsal medial PFC as well as the thalamus, right anterior insula and inferior frontal gyrus (IFG) (Figure 2C and Supplemental Table 4: Cluster 1). The second large cluster of positive FC was focused in the left PCC and spread into the left hippocampus (Cluster 4). Additional clusters of positive FC were found bilaterally in the angular gyrus (Cluster 8), posterior insula (Cluster 7), and superior temporal gyri (STG) (Cluster 7). One large cluster of negative FC was found in both hemispheres dorsally in the precuneus, SPL and occipital gyri (Cluster 2).

During exposure to the C-HP model the ICN derived from the bilateral pACC seed spread into the bilateral MPFC, superior frontal gyri (SFG), putamen and thalamus (Figure 2D and Supplemental Table 5: Cluster 1). The second cluster of positive FC encompassed the bilateral medial precuneus and PCC (Cluster 3). A third cluster of positive was found in the left DLPFC (Cluster 6). A large cluster of negative FC in the cerebellum spread bilaterally into the fusiform and MTG (Cluster 2). A second large cluster of negative FC spanned the right and left hemispheres and was focused in the right SPL and left IPL, angular and supramarginal gyri (Cluster 4).

The positive sACC FC cluster found during the non-pain scan spread into the left medial frontal gyrus left MFG and bilaterally in the SFG, as well as the right anterior insula (Figure 3A and Supplemental Table 6: Cluster 1). The second large cluster of positive FC was found in the MTG and STG (Cluster 5). Several clusters of negative FC included bilateral clusters in the MFG (Cluster 7), SFG (Cluster 8), as well as the right supramarginal gyrus and left IPL (Cluster 9).

**Figure 3.**
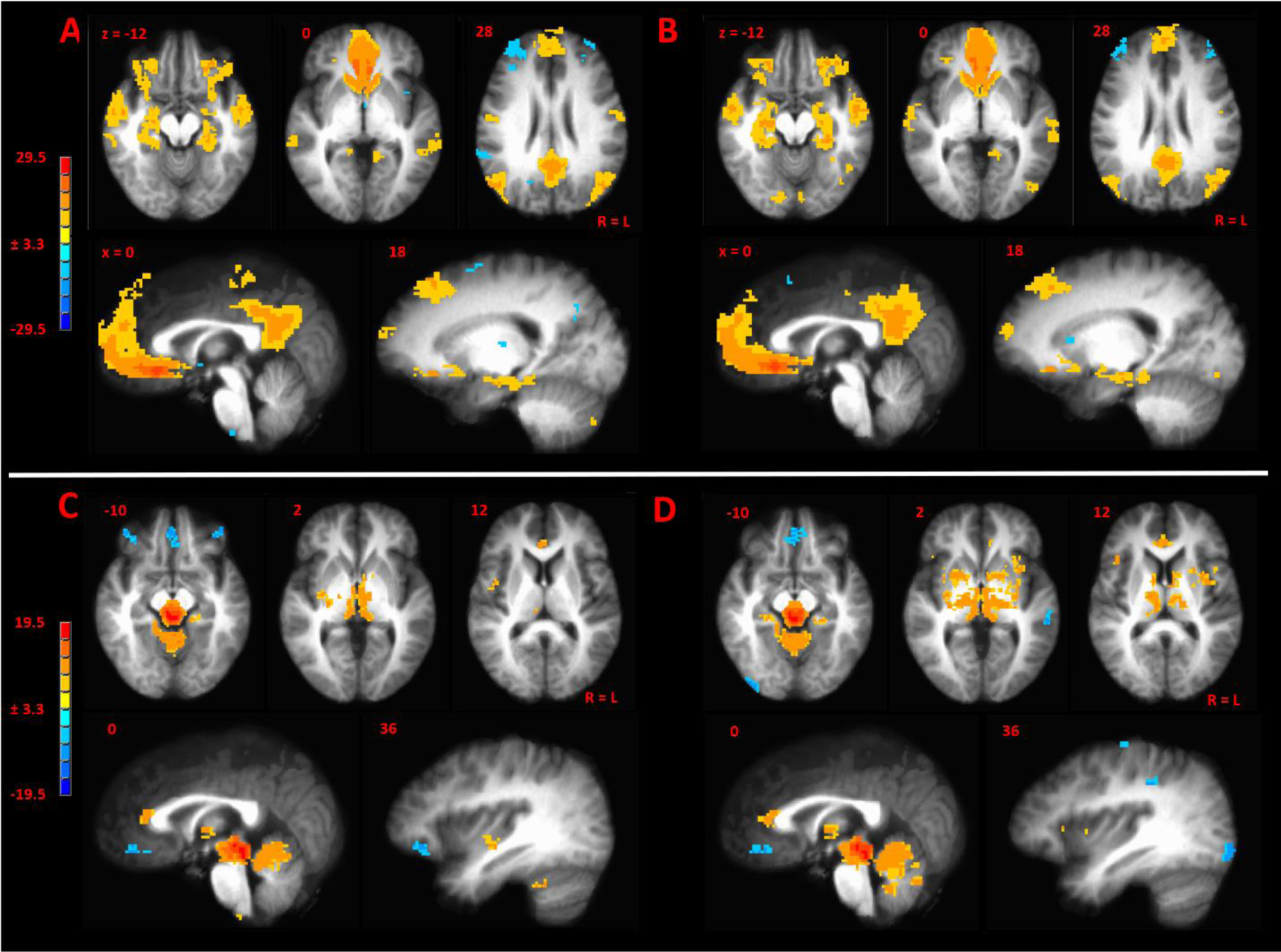
**A)** Mean seed-driven functional connectivity (SDFC) map from the sACC seed during the pain-free task. **B)** Mean SDFC map from the sACC seed during prolonged tonic pain. **C)** Mean SDFC map from the PAG seed during the pain-free task. **D)** Mean SDFC map from the PAG seed during prolonged tonic pain. R=L; Bar is in Z-units.

During exposure to the C-HP model the ICN derived from the bilateral sACC seed spread into the bilateral medial and SFG as well as the right anterior and middle insula (Figure 3B and Supplemental Table 7: Cluster 1). The second cluster of positive FC encompassed the bilateral PCC (Cluster 2). Several individual clusters on negative FC were found regions including the right MFG (Cluster 4) and right supramarginal gyrus (Cluster 5).

The control ICN map derived from the PAG seed included a large cluster of positive FC encompassing the midbrain as well as medial and anterior thalamus bilaterally (Figure 3C and Supplemental Table 8: Cluster 1). Clusters of positive FC was also found in the pACC (Cluster 3), right middle insula (Cluster 5), left PHG (Cluster 6) and a brainstem region consistent with the nucleus raphe magnus (Cluster 10). Clusters of negative FC were found in the right and left MFG (Clusters 2, 7 and 8). During prolonged tonic pain, the large cluster of positive FC emanating from the PAG encompassed the midbrain and spread dorsally and anteriorly into the medial thalamus bilaterally, bilateral caudate and putamen, right IFG and left anterior insula (Figure 3D and Supplemental Table 9: Cluster 1). Additional clusters of positive FC were found in the pACC (Cluster 2), left PHG (Cluster 9), and an area of the brainstem consistent with rostral ventral medulla (Cluster 17). Clusters of negative FC were found in the right inferior occipital gyrus (Cluster 3), left DLPFC (Cluster 5), left ventromedial prefrontal gyrus (Cluster 4), and right IPL (Clusters 11 and 13).

### Resting State Functional Connectivity: Brain-wide contrast between pain and pain-free conditions

To determine the statistically significant difference between the seed-derived FC of the resting state maps collected during the pain-free task and prolonged tonic pain, we calculated brain-wide contrast maps (Figure 4). The contrast map derived from the ICN maps of the bilateral aMCC revealed two significant clusters of enhanced FC during prolonged tonic pain in the left paracentral lobule (PCL) and left supplementary motor area (SMA) (Figure 4A and Table 1). The contrast map derived from the ICN maps of the bilateral pACC revealed significant clusters of enhanced FC during the prolonged tonic pain bilaterally in the IPL, right medial dorsal (MD) thalamus and right cerebellum (Figure 4B and Table 1). Additionally, the contrast of seed-driven FC from pACC showed reduced FC during the prolonged tonic pain in the left SFG. The contrast map derived from the ICN maps of the bilateral sACC revealed a significant cluster of enhanced FC during prolonged tonic pain in the left cerebellum (Figure 4C and Table 1). Additionally, the contrast of seed-driven FC from sACC showed reduced FC during prolonged tonic pain in two clusters in the right cerebellum and a cortical cluster in the right angular gyrus. The contrast map derived from the ICN maps of the PAG revealed a significant cluster of enhanced FC during prolonged tonic pain in the left lateral globus pallidus (Figure 4D and Table 1). The contrast of seed-driven FC from PAG showed reduced FC during prolonged tonic pain in a cluster in the right precentral gyrus and the left DLPFC. The DLPFC cluster resulted from positive FC between the PAG and the left DLPFC during the pain-free task changing to negative FC during prolonged tonic pain.

**Figure 4.**
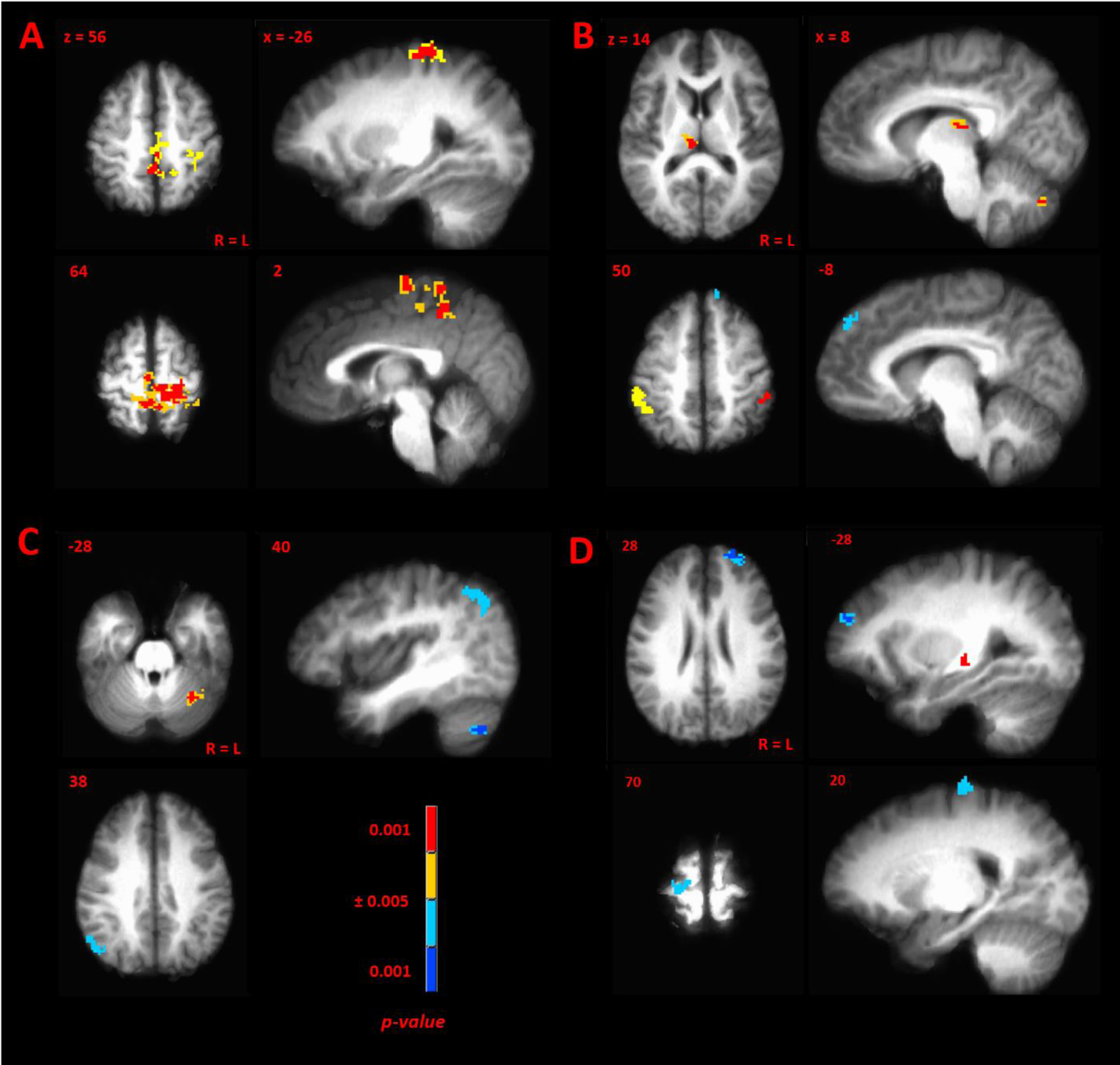
Contrast FC maps pain>control state for the following seeds: **A)** aMCC; **B)** pACC; **C)** sACC; **D)** PAG. P-value threshold was 0.005 with a cluster extent correction of 606 mm^3^ for cerebral cortex (Supplemental Table 1), R=L.

**Table 1.**
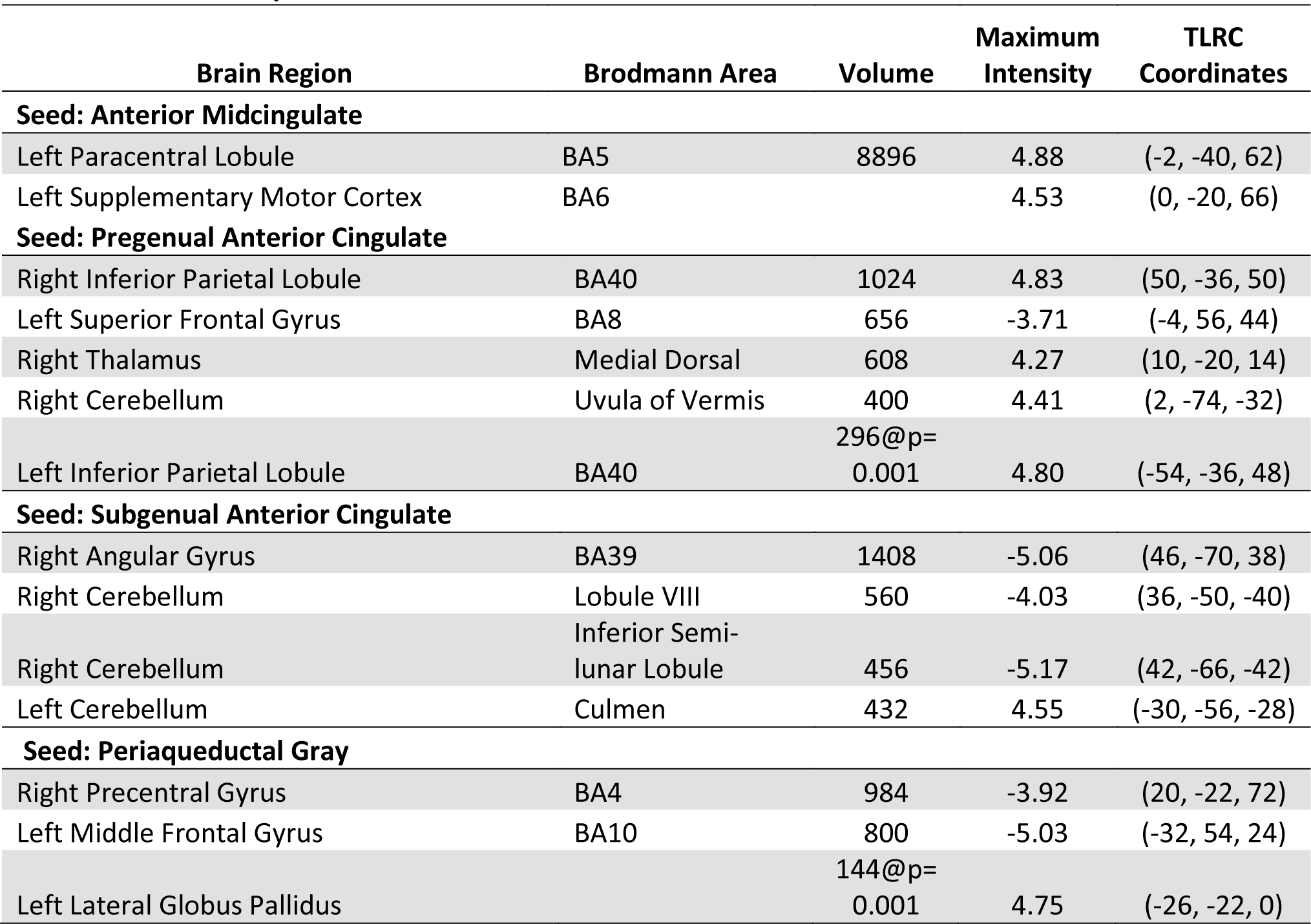
Contrast Maps Pain-Control.

| Brain Region | Brodmann Area | Volume | Maximum Intensity | TLRC Coordinates |
| --- | --- | --- | --- | --- |
| **Seed: Anterior Midcingulate** |  |  |  |  |
| Left Paracentral Lobule | BA5 | 8896 | 4.88 | (-2, -40, 62) |
| Left Supplementary Motor Cortex | BA6 |  | 4.53 | (0, -20, 66) |
| **Seed: Pregenua Anterior Cingulate** |  |  |  |  |
| Right Inferior Parietal Lobule | BA40 | 1024 | 4.83 | (50, -36, 50) |
| Left Superior Frontal Gyrus | BA8 | 656 | -3.71 | (-4, 56, 44) |
| Right Thalamus | Medial Dorsal | 608 | 4.27 | (10, -20, 14) |
| Right Cerebellum | Uvula of Vermis | 400 | 4.41 | (2, -74, -32) |
| Left Inferior Parietal Lobule | BA40 | 296@p=0.001 | 4.80 | (-54, -36, 48) |
| **Seed: Subgenual Anterior Cingulate** |  |  |  |  |
| Right Angular Gyrus | BA39 | 1408 | -5.06 | (46, -70, 38) |
| Right Cerebellum | Lobule VIII | 560 | -4.03 | (36, -50, -40) |
| Right Cerebellum | Inferior Semi-lunar Lobule | 456 | -5.17 | (42, -66, -42) |
| Left Cerebellum | Culmen | 432 | 4.55 | (-30, -56, -28) |
| **Seed: Periaqueductal Gray** |  |  |  |  |
| Right Precentral Gyrus | BA4 | 984 | -3.92 | (20, -22, 72) |
| Left Middle Frontal Gyrus | BA10 | 800 | -5.03 | (-32, 54, 24) |
| Left Lateral Globus Pallidus |  | 144@p=0.001 | 4.75 | (-26, -22, 0) |

### Resting State Functional Connectivity: Pain intensity covariance with seed-based FC

In the pain intensity covariation analysis (PPI analysis), greater pain intensity was related to weaker FC during prolonged tonic pain between the aMCC and clusters in the right IFG, left SFG, PCL, thalamus, putamen and posterior insula (Figure 5A and Table 2). Additionally, weaker FC during prolonged tonic pain between the pACC and clusters in the right precentral gyrus and the left SFG and caudate were related to greater pain intensity (Figure 5B and Table 2). Finally, stronger FC during prolonged tonic pain between the PAG and clusters in the bilateral SPL and left cerebellum were related to greater pain intensity (Figure 5C and Table 2).

**Figure 5.**
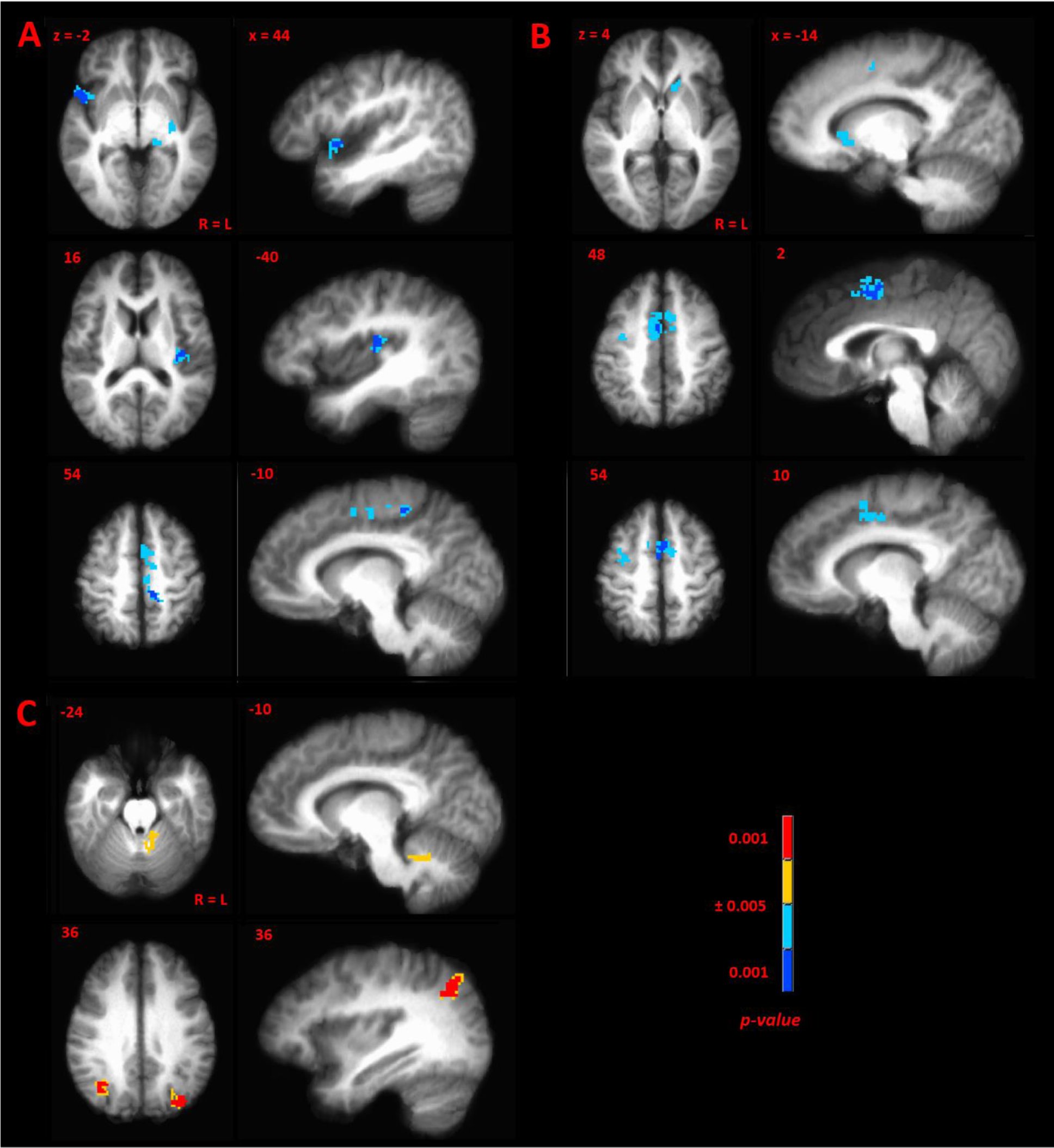
Pain intensity covariation during prolonged tonic pain for the following seeds: **A)** aMCC; **B)** pACC; **C)** PAG. P-value threshold was 0.005 with a cluster extent correction of 606 mm^3^ for cerebral cortex (Supplemental Table 1), R=L.

**Table 2.** Pain Intensity Covariation Maps

| Brain Region | Brodmann Area | Volume | Maximum Intensity | TLRC Coordinates |
| --- | --- | --- | --- | --- |
| **Seed: Anterior Midcingulate** |  |  |  |  |
| Right Inferior Frontal Gyrus | BA47 | 1192 | -4.68 | (50, 12, 0) |
| Left Superior Frontal Gyrus | BA6 | 840 | -4.17 | (-2, 4, 54) |
| Left Paracentral Lobule | BA5 | 816 | -4.42 | (-14, -36, 52) |
| Left Posterior Insula | BA13 | 784 | -4.89 | (-40, -20, 16) |
| Left Putamen |  | 336 | -4.78 | (-30, -12, -4) |
| Left Thalamus | Pulvinar | 328 | -4.68 | (-16, -28, 0) |
| **Seed: Pregenua Anterior Cingulate** |  |  |  |  |
| Left Superior Frontal Gyrus | BA6 | 3720 | -4.76 | (-2, 6, 50) |
| Right Precentral Gyrus | BA6 | 624 | -4.26 | (32, -8, 50) |
| Left Caudate |  | 456 | -4.06 | (-14, 22, 6) |
| **Seed: Periaqueductal Gray** |  |  |  |  |
| Right Precuneus/Superior Parietal Lobule | BA19/BA7 | 1968 | 4.70 | (34, -66, 36) |
| Left Precuneus/Superior Parietal Lobule | BA19/BA7 | 1152 | 5.97 | (-34, -78, 36) |
| Left Cerebellum | Fastigium/Dentate | 424 | 4.14 | (-8, -52, -24) |

The sACC seed-driven pain intensity-FC covariation maps during prolonged tonic pain showed no significant clusters and the global minimum q-value was 0.99. Furthermore, all seed-driven pain intensity-FC covariation maps during the control state, using the pain intensities reported during the C-HP model, had q-values indicative of few true voxel detections (aMCC, q=1.00; pACC, q=0.37; sACC, q=1.00; PAG, q=0.41), indicating that the relationship between pain intensity and DPMN FC is dependent on the prolonged tonic painful stimuli.

## Discussion

We evaluated the role of prolonged tonic pain in modulating resting state FC of the descending pain modulatory network (DPMN). While our predictions of diminished connectivity to DPMN seeds was upheld, observed modulations of FC associated with prolonged tonic pain were found in structures outside our hypothesized network. Specifically, we found enhanced FC between aMCC and left PCL somatosensory foot representation (Figure 4A). Additionally, we found enhanced FC during prolonged tonic pain between pACC and bilateral IPL and right MD thalamus, with reduced FC between left DLPFC and pACC (Figure 4B). FC between bilateral sACC and right angular gyrus was reduced during prolonged tonic pain. PAG FC during prolonged tonic pain was characterized by disruptions with precentral gyrus and left medial frontal gyrus (Figure 4D). In general, DPMN nodes showed a complex pattern of FC enhancements and disruptions with brain regions linked to sensory discrimination, affective and motivational aspects of pain-related neural processing [18;47;60;74;107].

### Prolonged pain-driven reversal of RSFC between L-DLPFC and PAG

Our PAG-seeded ICN resembled previously reported FC maps. We found FC peaks in bilateral cerebellum, bilateral pACC and sACC, left DLPFC and left anterior thalamus. This distribution of FC is consistent with the known axonal connections of the PAG [1;37;69;90]. Previously reported PAG FC maps in healthy subjects demonstrated significant FC with bilateral cerebellum, pACC, medial and anterior thalamus [45;51;61;65;94;108].

We observed positive FC between PAG and left DLPFC during the pain-free state, consistent with white matter tractography and treatment-induced augmentation of FC between the PAG and DLPFC in pediatric complex regional pain syndrome [23;30;37;88-90]. Previously reported PAG FC maps have some notable differences from our map. For example, unlike some previous maps, our map lacks any significant above threshold anticorrelated regions [45;108]. This is possibly because we avoided introducing spurious anticorrelations by not removing global signal or signal from grey matter in our WM and CSF masks [14;79]. Additionally, many maps included regions of the pontine brainstem and rostral ventral medulla [45;94;108]. This discrepancy may be caused by our use of a strict cluster threshold designed to minimize corresponding voxel- and brain-wise FDR [19;29;104].

Prolonged tonic pain was associated with reduced FC between PAG and left DLPFC and between the pACC and left DLPFC, although in non-overlapping areas. These findings are consistent with results from several populations of chronic pain patients and activation changes in response to thermal allodynia in studies of capsaicin sensitization [59]. Reduced FC between the PAG and left DLPFC has been reported in migraine patients and women with dysmenorrhea [16;61;82;102]. Importantly, treatment-induced enhancement of FC in pediatric CRPS patients, enhanced coactivation and corelease of endogenous opioids in both the PAG and DLPFC during the analgesic period after invasive motor cortex stimulation, and enhanced FC of PAG and left DLPFC during placebo analgesia has been reported [23;30;58;72;97]. Several reports demonstrate many chronic pain disorders lead to gray matter or cortical thickness reductions in left DLPFC and PAG, and these reductions normalize after pain-alleviating treatment [12;22;88;89]. This is the first report to reveal PAG FC with the left DLPFC and reversal of this positive FC during experience of tonic prolonged pain.

### Seed-driven pACC and aMCC maps and the effects of prolonged tonic pain

The bilateral pACC and aMCC ICN maps were remarkably similar to the corresponding mean FC maps previously reported from these seed regions [63;83;92]. Each network, and their primary nodes, has been implicated in a variety of psychological processes, from conflict monitoring and mediation of top-down attention to the sense of oneself from which a unified feeling of consciousness emerges [10;20;24]. The aMCC and associated network appear involved in the continuous environmental monitoring, using ecological context to weigh incoming stimuli allowing a dynamic decision regarding stimulus saliency [8;75;76;98;103].

In the pACC ICN contrast, we found enhanced FC during prolonged tonic pain between the right MD thalamus and the aMCC seed. This connection is frequently seen in coactivation in response to acute pain and during the retention of memories of painful stimuli compared to pain experience [26;95]. Furthermore, patients with painful diabetic neuropathy show enhanced FC of PAG with MD thalamus and pACC, while patients with irritable bowel syndrome demonstrate reduced FC between MD thalamus and pACC during painful bowel distention [56;84]. Indeed, thalamocortical asynchrony is an hypothesized driver of persistent neuropathic pain, evidence from human and animal studies supports this hypothesis and recent evidence extends this mechanism to prolonged tonic pain from the C-HP model [32;39;40;57;80;81;87]. Our results further support a modulation of a specific connection between the pACC and MD and other midline thalamic nuclei [99;100].

The aMCC demonstrated enhanced FC with left PCL during C-HP stimulation of the leg. Enhanced FC between the aMCC and somatomotor cortex has been previously reported during painful pressure in healthy subjects [44]. Chronic pain patients including those with fibromyalgia, migraine, cluster headache, and primary dysmenorrhea also demonstrate enhanced FC between the aMCC and somatomotor cortex compared to healthy controls [27;43;55]. This finding may be particularly important as enhancement of FC between aMCC and somatosensory representation of the body part affected by painful stimuli is correlated with pain intensity, whether experimental or clinical ([43], Figure 5A).

### Pain intensity correlation with FC of DPMN

We found significant anticorrelations between pain intensity and FC during prolonged tonic pain between aMCC and left PCL, posterior insula, and thalamus (Figure 5A). The aMCC, PCL, and posterior insula are commonly activated by painful stimuli and neural responses in these regions are positively graded with subjective pain intensity [3;9;18;26;73;74;99;101]. Since FC is driven by correlations in BOLD signal, it may seem surprising that areas normally responsive to painful stimuli are anticorrelated. However, the BOLD response is known to vary in its temporal relationship to discrete stimuli depending on where the cortical response originates [38]. Therefore, a tonic stimulus varying in perceptual intensity over minutes would likely create BOLD signals more uncorrelated over time. We found negative correlation of pain intensity with FC between pACC and a bilateral cluster in the SMA and MCC (Figure 5B). The pACC is another region responsive to painful stimuli [26]. The negative correlation found here may reflect a similar process to aMCC FC relationships with pain intensity.

In contrast, pain intensity positively correlated with FC between PAG and bilateral SPL (Figure 5C). SPL is particularly important in painful stimuli discrimination by intensity and location [68;106]. Both the PAG and SPL bilaterally show enhanced BOLD responses to brush allodynia after capsaicin-induced sensitization in healthy subjects [62]. Central sensitization of spinal dorsal horn (SDH) neurons is mediated by an increase in excitatory amino acid receptor responses to excitatory amino acid release in the SDH and a reduction in the effectiveness of inhibitory amino acid mediated suppression of SDH activity in response to noxious stimuli [25;50]. After induction of central sensitization in SDH by intradermal injection of capsaicin in monkeys, descending inhibition from PAG is suppressed at the SDH. Therefore, in healthy subjects during primary afferent barrage resulting in central sensitization, the PAG likely increases its activity to compensate for the reduced effectiveness of inhibition at the SDH [50;64]. In concert with enhanced activation of the PAG, central sensitization by capsaicin enhances SPL and IPL responses to mechanical stimuli; and inhibitory repetitive transcranial magnetic stimulation of the parietal cortex inhibits secondary hyperalgesia after capsaicin exposure [48;59;85]. Therefore, our finding of positive correlation of pain intensity with enhanced PAG FC with bilateral SPL possibly results from the PAG providing compensatory inhibition of noxious input and the SPL processing expansion and enhancement of nociceptive gain during central sensitization.

### Limitations

Some methodological choices may limit interpretation of our results. For example, we choose not to conduct our analyses with global signal regression or motion scrubbing [77;79]. Regarding motion correction we made this choice because we felt our motion parameters were sufficiently stringent. We felt that since our analysis controlled for estimated motion parameters and their first differentials, this was sufficient to limit the effect of magnetic field distortion. Not using global signal regression is consistent with our regressors of no interest sufficiently controlled for CSF and WM noise, and avoiding spurious anticorrelation in our FC maps. BOLD signal is inherently supergaussian and global signal regression distorts this distribution [11;79].

### Conclusions

We report the effects of prolonged tonic pain generated by the capsaicin-heat pain model on descending pain modulatory network (DPMN) functional connectivity (FC). We found evidence supporting the DLPFC in modulating activity of the PAG and enhanced entrainment of SPL and PAG activity with increasing subjective pain report. We report enhanced FC of aMCC with PCL, coupled with a negative correlation of pain intensity with FC between aMCC and PCL during ongoing pain. Finally, FC of pACC with MD thalamus was enhanced during prolonged tonic pain. These results are consistent with several findings in chronic pain populations indicating that these previous findings of modulated FC in patients may be attributed to ongoing pain. Further studies contrasting FC in pain patients with well-matched controls experiencing prolonged tonic pain are needed to disentangle ongoing pain-related neural processing from disease-specific effects in chronic pain disorders.

## Acknowledgements

This work was supported by NIH grant P30-NR014019 and the University of Maryland Center to Advance Chronic Pain Research. TJM acknowledges support from the Johns Hopkins Neurosurgical Pain Research Institute and NIH grants R56-NS038493 and R01-NS107602. We gratefully acknowledge the technical assistance of Hee Jun Kim, Brooks DuBose, and Sean M. Cooper. All authors declare no conflicts of interest.

---

**Supplemental Figure 1.**
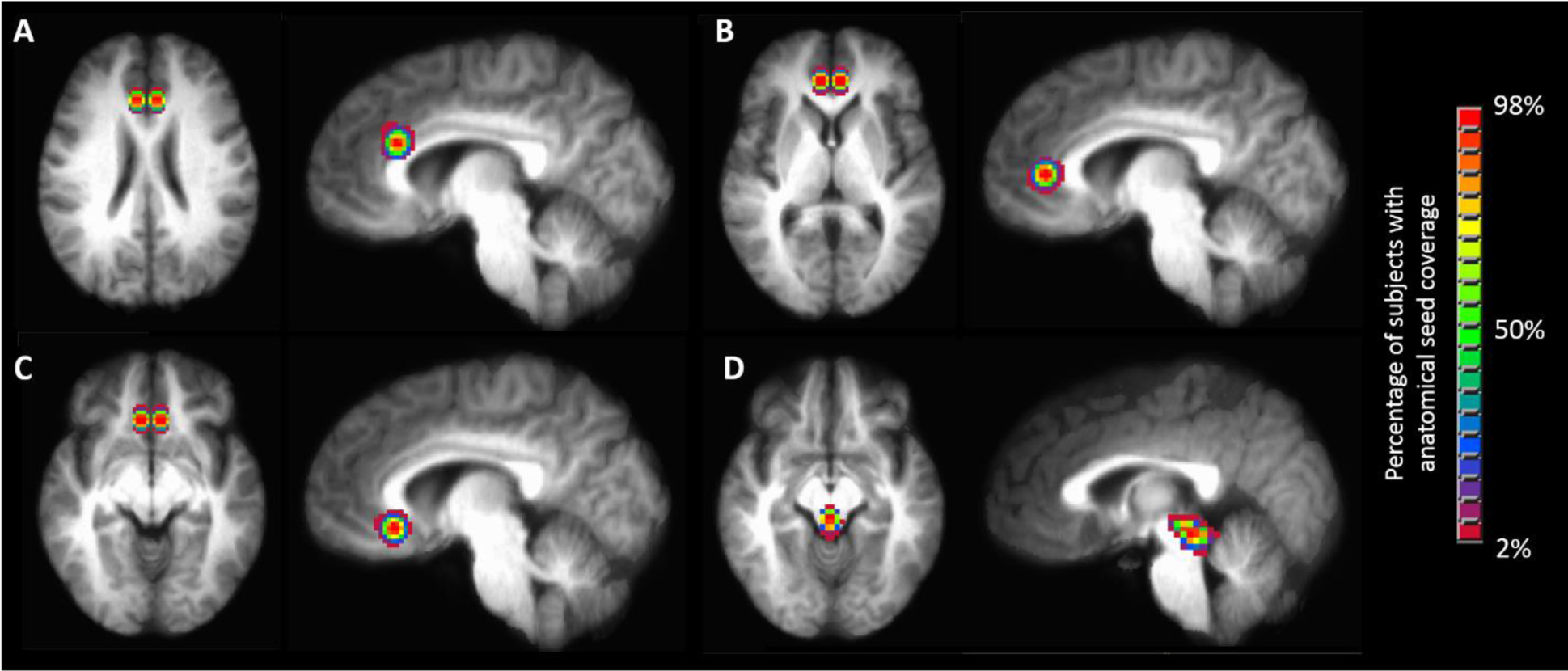
**A)** aMCC seed. **B)** pACC seed. **C)** sACC seed. **D)** PAG seed.

**Supplemental Figure 2.**
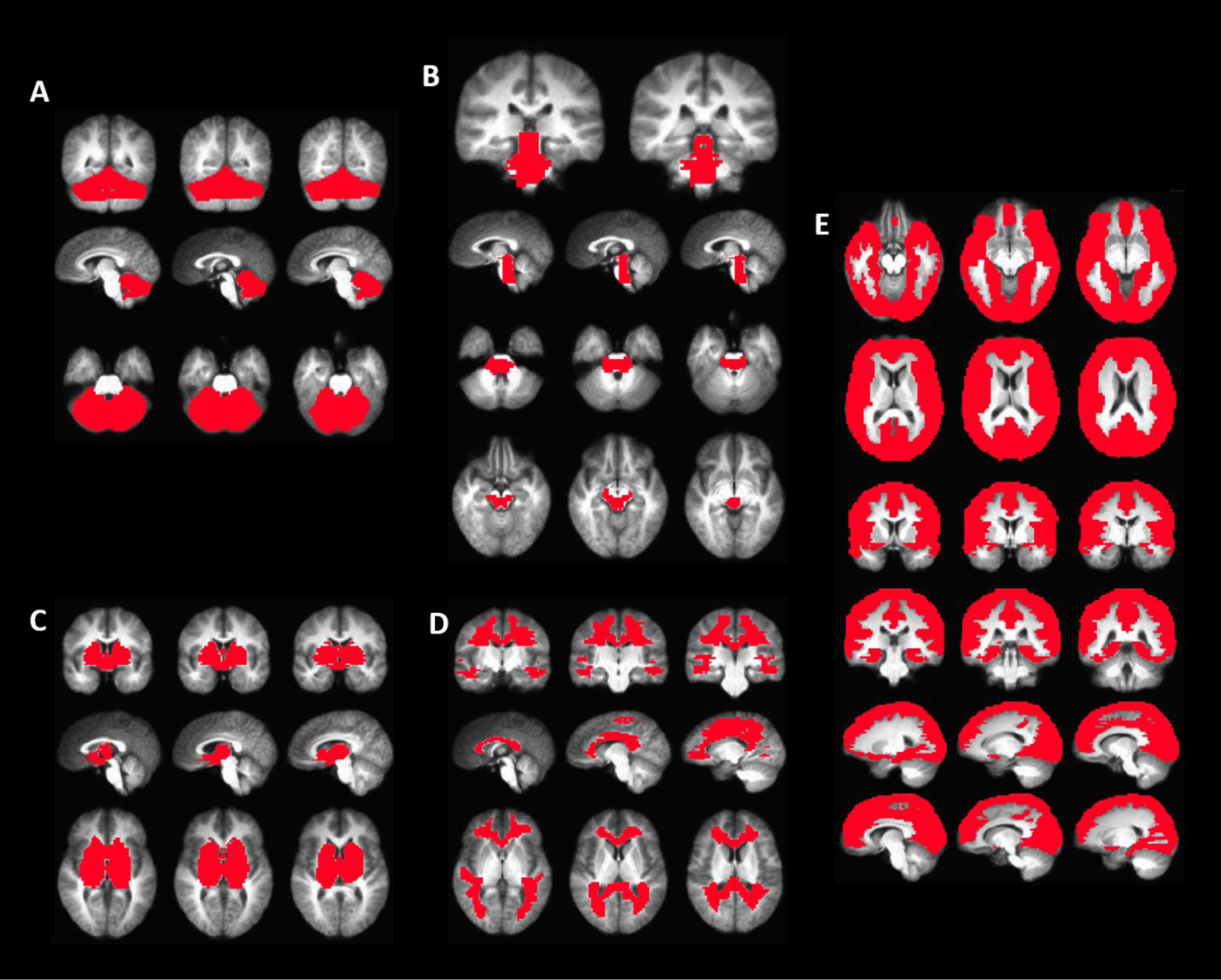
**A)** Cerebellum functional mask. **B)** Brainstem functional mask. **C)** Subcortical gray matter mask. **D)** White matter mask. **E)** Cortical functional mask.

**Supplemental Table 1.** Cluster volume for the appropriate Cluster Extent Criteria for each p-value level shown

| **Anatomical Mask** | **p-value** | **Volume (mm<sup>3</sup>)** |
| --- | --- | --- |
| **Cortical** | 0.005 | 606 |
|  | 0.001 | 253 |
| **Cerebellar** | 0.005 | 334 |
|  | 0.001 | 139 |
| **Subcortical Gray Matter** | 0.005 | 288 |
|  | 0.001 | 120 |
| **Brainstem** | 0.005 | 160 |
|  | 0.001 | 64 |

**Supplemental Table 2.**
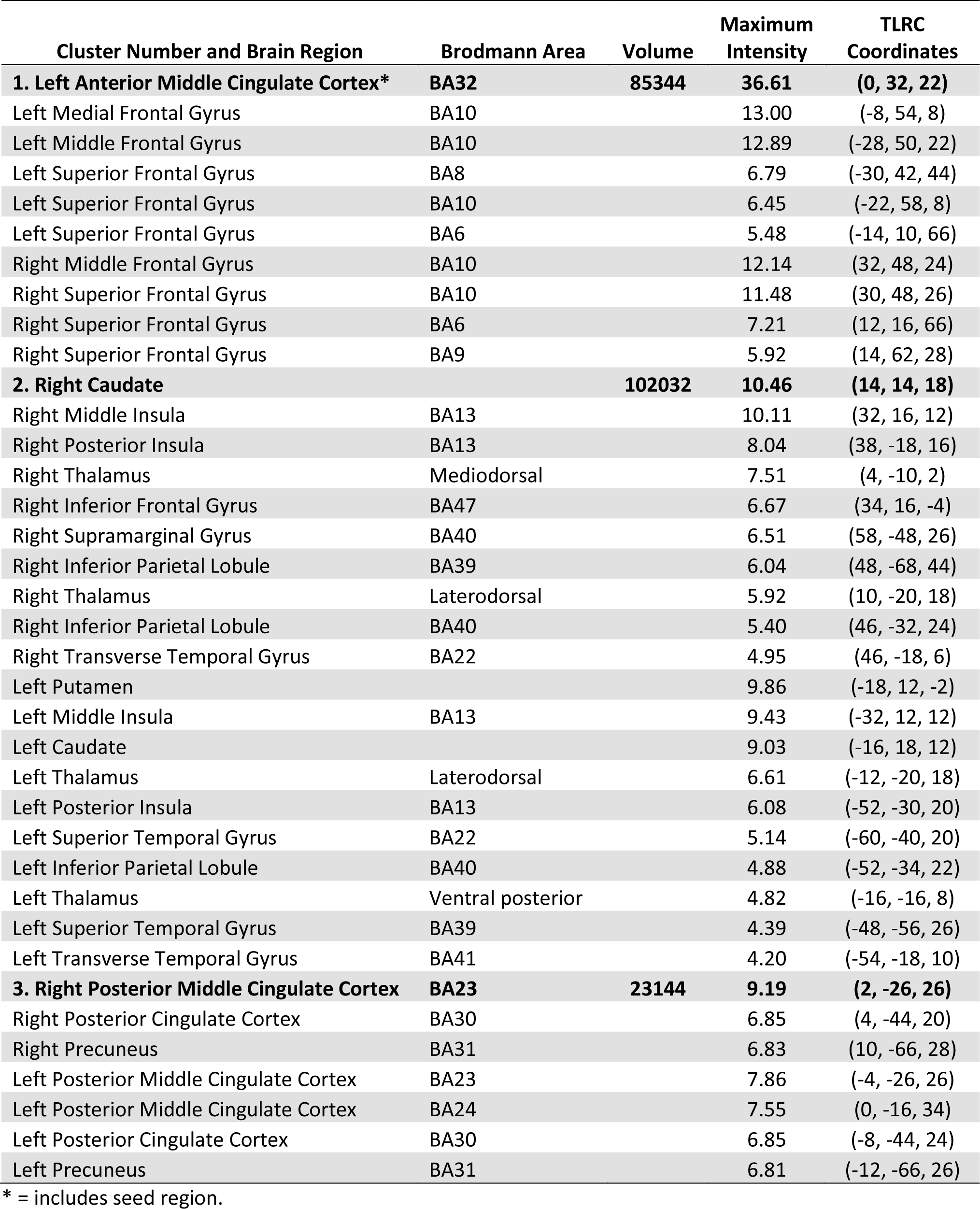
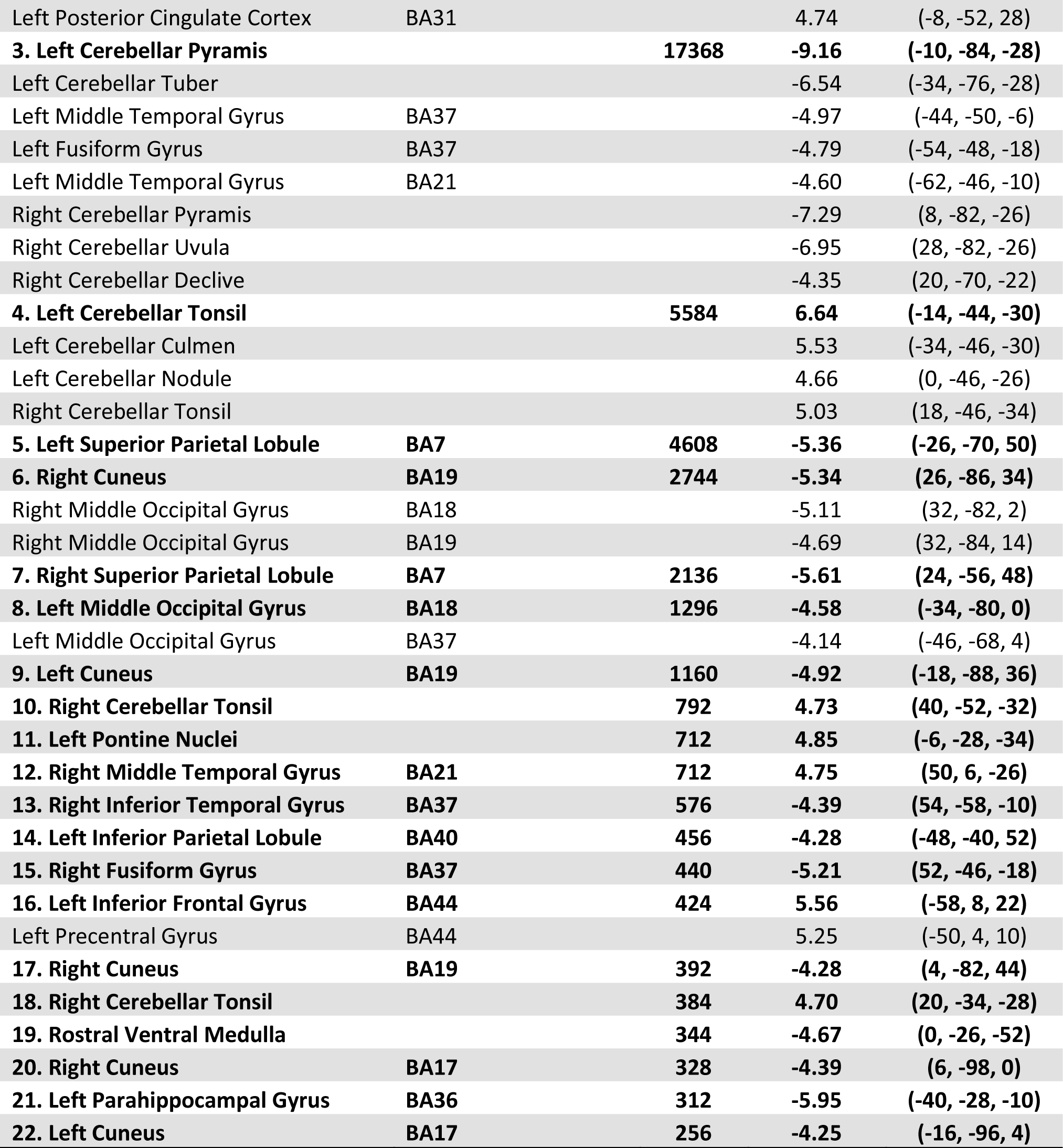
Seed: Bilateral Anterior MCC State: Control

| Cluster Number and Brain Region | Brodmann Area | Volume | Maximum Intensity | TLRC Coordinates |
| --- | --- | --- | --- | --- |
| **1. Left Anterior Middle Cingulate Cortex\*** | **BA32** | **85344** | **36.61** | **(0, 32, 22)** |
| Left Medial Frontal Gyrus | BA10 |  | 13.00 | (-8, 54, 8) |
| Left Middle Frontal Gyrus | BA10 |  | 12.89 | (-28, 50, 22) |
| Left Superior Frontal Gyrus | BA8 |  | 6.79 | (-30, 42, 44) |
| Left Superior Frontal Gyrus | BA10 |  | 6.45 | (-22, 58, 8) |
| Left Superior Frontal Gyrus | BA6 |  | 5.48 | (-14, 10, 66) |
| Right Middle Frontal Gyrus | BA10 |  | 12.14 | (32, 48, 24) |
| Right Superior Frontal Gyrus | BA10 |  | 11.48 | (30, 48, 26) |
| Right Superior Frontal Gyrus | BA6 |  | 7.21 | (12, 16, 66) |
| Right Superior Frontal Gyrus | BA9 |  | 5.92 | (14, 62, 28) |
| **2. Right Caudate** |  | **102032** | **10.46** | **(14, 14, 18)** |
| Right Middle Insula | BA13 |  | 10.11 | (32, 16, 12) |
| Right Posterior Insula | BA13 |  | 8.04 | (38, -18, 16) |
| Right Thalamus | Mediodorsal |  | 7.51 | (4, -10, 2) |
| Right Inferior Frontal Gyrus | BA47 |  | 6.67 | (34, 16, -4) |
| Right Supramarginal Gyrus | BA40 |  | 6.51 | (58, -48, 26) |
| Right Inferior Parietal Lobule | BA39 |  | 6.04 | (48, -68, 44) |
| Right Thalamus | Laterodorsal |  | 5.92 | (10, -20, 18) |
| Right Inferior Parietal Lobule | BA40 |  | 5.40 | (46, -32, 24) |
| Right Transverse Temporal Gyrus | BA22 |  | 4.95 | (46, -18, 6) |
| Left Putamen |  |  | 9.86 | (-18, 12, -2) |
| Left Middle Insula | BA13 |  | 9.43 | (-32, 12, 12) |
| Left Caudate |  |  | 9.03 | (-16, 18, 12) |
| Left Thalamus | Laterodorsal |  | 6.61 | (-12, -20, 18) |
| Left Posterior Insula | BA13 |  | 6.08 | (-52, -30, 20) |
| Left Superior Temporal Gyrus | BA22 |  | 5.14 | (-60, -40, 20) |
| Left Inferior Parietal Lobule | BA40 |  | 4.88 | (-52, -34, 22) |
| Left Thalamus | Ventral posterior |  | 4.82 | (-16, -16, 8) |
| Left Superior Temporal Gyrus | BA39 |  | 4.39 | (-48, -56, 26) |
| Left Transverse Temporal Gyrus | BA41 |  | 4.20 | (-54, -18, 10) |
| **3. Right Posterior Middle Cingulate Cortex** | **BA23** | **23144** | **9.19** | **(2, -26, 26)** |
| Right Posterior Cingulate Cortex | BA30 |  | 6.85 | (4, -44, 20) |
| Right Precuneus | BA31 |  | 6.83 | (10, -66, 28) |
| Left Posterior Middle Cingulate Cortex | BA23 |  | 7.86 | (-4, -26, 26) |
| Left Posterior Middle Cingulate Cortex | BA24 |  | 7.55 | (0, -16, 34) |
| Left Posterior Cingulate Cortex | BA30 |  | 6.85 | (-8, -44, 24) |
| Left Precuneus | BA31 |  | 6.81 | (-12, -66, 26) |

\* = includes seed region.

| Cluster Number and Brain Region | Brodmann Area | Volume | Maximum Intensity | TLRC Coordinates |
| --- | --- | --- | --- | --- |
| Left Posterior Cingulate Cortex | BA31 |  | 4.74 | (-8, -52, 28) |
| **3. Left Cerebellar Pyramis** |  | **17368** | **-9.16** | **(-10, -84, -28)** |
| Left Cerebellar Tuber |  |  | -6.54 | (-34, -76, -28) |
| Left Middle Temporal Gyrus | BA37 |  | -4.97 | (-44, -50, -6) |
| Left Fusiform Gyrus | BA37 |  | -4.79 | (-54, -48, -18) |
| Left Middle Temporal Gyrus | BA21 |  | -4.60 | (-62, -46, -10) |
| Right Cerebellar Pyramis |  |  | -7.29 | (8, -82, -26) |
| Right Cerebellar Uvula |  |  | -6.95 | (28, -82, -26) |
| Right Cerebellar Declive |  |  | -4.35 | (20, -70, -22) |
| **4. Left Cerebellar Tonsil** |  | **5584** | **6.64** | **(-14, -44, -30)** |
| Left Cerebellar Culmen |  |  | 5.53 | (-34, -46, -30) |
| Left Cerebellar Nodule |  |  | 4.66 | (0, -46, -26) |
| Right Cerebellar Tonsil |  |  | 5.03 | (18, -46, -34) |
| **5. Left Superior Parietal Lobule** | **BA7** | **4608** | **-5.36** | **(-26, -70, 50)** |
| **6. Right Cuneus** | **BA19** | **2744** | **-5.34** | **(26, -86, 34)** |
| Right Middle Occipital Gyrus | BA18 |  | -5.11 | (32, -82, 2) |
| Right Middle Occipital Gyrus | BA19 |  | -4.69 | (32, -84, 14) |
| **7. Right Superior Parietal Lobule** | **BA7** | **2136** | **-5.61** | **(24, -56, 48)** |
| **8. Left Middle Occipital Gyrus** | **BA18** | **1296** | **-4.58** | **(-34, -80, 0)** |
| Left Middle Occipital Gyrus | BA37 |  | -4.14 | (-46, -68, 4) |
| **9. Left Cuneus** | **BA19** | **1160** | **-4.92** | **(-18, -88, 36)** |
| **10. Right Cerebellar Tonsil** |  | **792** | **4.73** | **(40, -52, -32)** |
| **11. Left Pontine Nuclei** |  | **712** | **4.85** | **(-6, -28, -34)** |
| **12. Right Middle Temporal Gyrus** | **BA21** | **712** | **4.75** | **(50, 6, -26)** |
| **13. Right Inferior Temporal Gyrus** | **BA37** | **576** | **-4.39** | **(54, -58, -10)** |
| **14. Left Inferior Parietal Lobule** | **BA40** | **456** | **-4.28** | **(-48, -40, 52)** |
| **15. Right Fusiform Gyrus** | **BA37** | **440** | **-5.21** | **(52, -46, -18)** |
| **16. Left Inferior Frontal Gyrus** | **BA44** | **424** | **5.56** | **(-58, 8, 22)** |
| Left Precentral Gyrus | BA44 |  | 5.25 | (-50, 4, 10) |
| **17. Right Cuneus** | **BA19** | **392** | **-4.28** | **(4, -82, 44)** |
| **18. Right Cerebellar Tonsil** |  | **384** | **4.70** | **(20, -34, -28)** |
| **19. Rostral Ventral Medulla** |  | **344** | **-4.67** | **(0, -26, -52)** |
| **20. Right Cuneus** | **BA17** | **328** | **-4.39** | **(6, -98, 0)** |
| **21. Left Parahippocampal Gyrus** | **BA36** | **312** | **-5.95** | **(-40, -28, -10)** |
| **22. Left Cuneus** | **BA17** | **256** | **-4.25** | **(-16, -96, 4)** |

**Supplemental Table 3.**
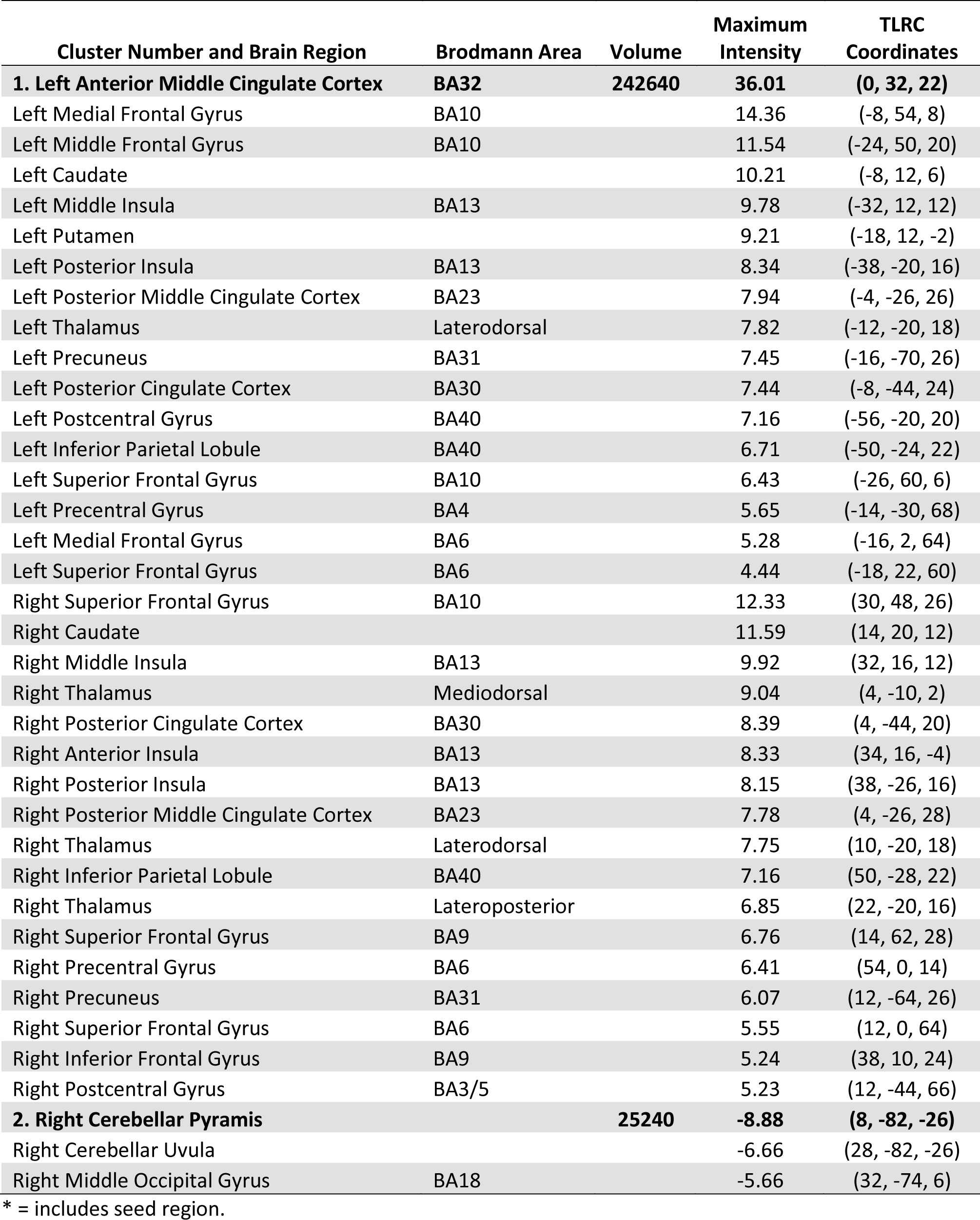
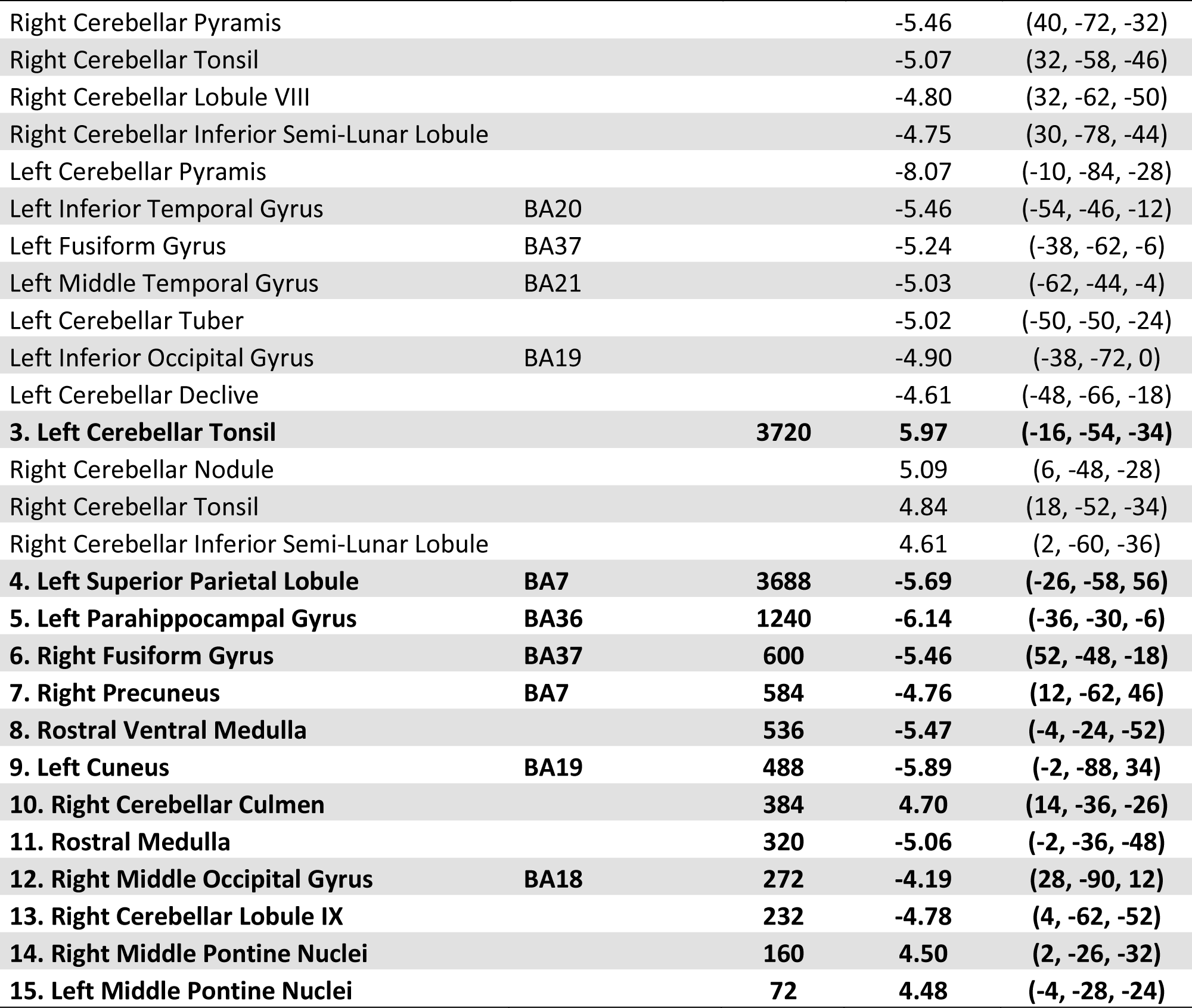
Seed: Bilateral Anterior MCC State: Pain

| Cluster Number and Brain Region | Brodmann Area | Volume | Maximum Intensity | TLRC Coordinates |
| --- | --- | --- | --- | --- |
| **1. Left Anterior Middle Cingulate Cortex** | **BA32** | **242640** | **36.01** | **(0, 32, 22)** |
| Left Medial Frontal Gyrus | BA10 |  | 14.36 | (-8, 54, 8) |
| Left Middle Frontal Gyrus | BA10 |  | 11.54 | (-24, 50, 20) |
| Left Caudate |  |  | 10.21 | (-8, 12, 6) |
| Left Middle Insula | BA13 |  | 9.78 | (-32, 12, 12) |
| Left Putamen |  |  | 9.21 | (-18, 12, -2) |
| Left Posterior Insula | BA13 |  | 8.34 | (-38, -20, 16) |
| Left Posterior Middle Cingulate Cortex | BA23 |  | 7.94 | (-4, -26, 26) |
| Left Thalamus | Laterodorsal |  | 7.82 | (-12, -20, 18) |
| Left Precuneus | BA31 |  | 7.45 | (-16, -70, 26) |
| Left Posterior Cingulate Cortex | BA30 |  | 7.44 | (-8, -44, 24) |
| Left Postcentral Gyrus | BA40 |  | 7.16 | (-56, -20, 20) |
| Left Inferior Parietal Lobule | BA40 |  | 6.71 | (-50, -24, 22) |
| Left Superior Frontal Gyrus | BA10 |  | 6.43 | (-26, 60, 6) |
| Left Precentral Gyrus | BA4 |  | 5.65 | (-14, -30, 68) |
| Left Medial Frontal Gyrus | BA6 |  | 5.28 | (-16, 2, 64) |
| Left Superior Frontal Gyrus | BA6 |  | 4.44 | (-18, 22, 60) |
| Right Superior Frontal Gyrus | BA10 |  | 12.33 | (30, 48, 26) |
| Right Caudate |  |  | 11.59 | (14, 20, 12) |
| Right Middle Insula | BA13 |  | 9.92 | (32, 16, 12) |
| Right Thalamus | Mediodorsal |  | 9.04 | (4, -10, 2) |
| Right Posterior Cingulate Cortex | BA30 |  | 8.39 | (4, -44, 20) |
| Right Anterior Insula | BA13 |  | 8.33 | (34, 16, -4) |
| Right Posterior Insula | BA13 |  | 8.15 | (38, -26, 16) |
| Right Posterior Middle Cingulate Cortex | BA23 |  | 7.78 | (4, -26, 28) |
| Right Thalamus | Laterodorsal |  | 7.75 | (10, -20, 18) |
| Right Inferior Parietal Lobule | BA40 |  | 7.16 | (50, -28, 22) |
| Right Thalamus | Lateroposterior |  | 6.85 | (22, -20, 16) |
| Right Superior Frontal Gyrus | BA9 |  | 6.76 | (14, 62, 28) |
| Right Precentral Gyrus | BA6 |  | 6.41 | (54, 0, 14) |
| Right Precuneus | BA31 |  | 6.07 | (12, -64, 26) |
| Right Superior Frontal Gyrus | BA6 |  | 5.55 | (12, 0, 64) |
| Right Inferior Frontal Gyrus | BA9 |  | 5.24 | (38, 10, 24) |
| Right Postcentral Gyrus | BA3/5 |  | 5.23 | (12, -44, 66) |
| **2. Right Cerebellar Pyramid** |  | **25240** | **-8.88** | **(8, -82, -26)** |
| Right Cerebellar Uvula |  |  | -6.66 | (28, -82, -26) |
| Right Middle Occipital Gyrus | BA18 |  | -5.66 | (32, -74, 6) |

\* = includes seed region.

| Cluster Number and Brain Region | Brodmann Area | Volume | Maximum Intensity | TLRC Coordinates |
| --- | --- | --- | --- | --- |
| Right Cerebellar Pyramis |  |  | -5.46 | (40, -72, -32) |
| Right Cerebellar Tonsil |  |  | -5.07 | (32, -58, -46) |
| Right Cerebellar Lobule VIII |  |  | -4.80 | (32, -62, -50) |
| Right Cerebellar Inferior Semi-Lunar Lobule |  |  | -4.75 | (30, -78, -44) |
| Left Cerebellar Pyramis |  |  | -8.07 | (-10, -84, -28) |
| Left Inferior Temporal Gyrus | BA20 |  | -5.46 | (-54, -46, -12) |
| Left Fusiform Gyrus | BA37 |  | -5.24 | (-38, -62, -6) |
| Left Middle Temporal Gyrus | BA21 |  | -5.03 | (-62, -44, -4) |
| Left Cerebellar Tuber |  |  | -5.02 | (-50, -50, -24) |
| Left Inferior Occipital Gyrus | BA19 |  | -4.90 | (-38, -72, 0) |
| Left Cerebellar Declive |  |  | -4.61 | (-48, -66, -18) |
| **3. Left Cerebellar Tonsil** |  | **3720** | **5.97** | **(-16, -54, -34)** |
| Right Cerebellar Nodule |  |  | 5.09 | (6, -48, -28) |
| Right Cerebellar Tonsil |  |  | 4.84 | (18, -52, -34) |
| Right Cerebellar Inferior Semi-Lunar Lobule |  |  | 4.61 | (2, -60, -36) |
| **4. Left Superior Parietal Lobule** | **BA7** | **3688** | **-5.69** | **(-26, -58, 56)** |
| **5. Left Parahippocampal Gyrus** | **BA36** | **1240** | **-6.14** | **(-36, -30, -6)** |
| **6. Right Fusiform Gyrus** | **BA37** | **600** | **-5.46** | **(52, -48, -18)** |
| **7. Right Precuneus** | **BA7** | **584** | **-4.76** | **(12, -62, 46)** |
| **8. Rostral Ventral Medulla** |  | **536** | **-5.47** | **(-4, -24, -52)** |
| **9. Left Cuneus** | **BA19** | **488** | **-5.89** | **(-2, -88, 34)** |
| **10. Right Cerebellar Culmen** |  | **384** | **4.70** | **(14, -36, -26)** |
| **11. Rostral Medulla** |  | **320** | **-5.06** | **(-2, -36, -48)** |
| **12. Right Middle Occipital Gyrus** | **BA18** | **272** | **-4.19** | **(28, -90, 12)** |
| **13. Right Cerebellar Lobule IX** |  | **232** | **-4.78** | **(4, -62, -52)** |
| **14. Right Middle Pontine Nuclei** |  | **160** | **4.50** | **(2, -26, -32)** |
| **15. Left Middle Pontine Nuclei** |  | **72** | **4.48** | **(-4, -28, -24)** |

**Supplemental Table 4.** Seed: Bilateral Pregenual ACC State: Control

| Cluster Number and Brain Region | Brodmann Area | Volume | Maximum Intensity | TLRC Coordinates |
| --- | --- | --- | --- | --- |
| **1. Left Pregenual Anterior Cingulate Cortex** | **BA32** | **114792** | **35.44** | **(-4, 40, 6)** |
| Left Caudate |  |  | 11.24 | (-14, 18, 6) |
| Left Putamen |  |  | 8.88 | (-20, 14, -4) |
| Left Thalamus | Mediodorsal |  | 6.48 | (0, -14, 8) |
| Left Superior Frontal Gyrus | BA8 |  | 5.65 | (-36, 16, 48) |
| Right Pregenual Anterior Cingulate Cortex | BA32 |  | 34.17 | (6, 42, 8) |
| Right Caudate |  |  | 12.91 | (14, 20, 4) |
| Right Superior Frontal Gyrus | BA9 |  | 9.96 | (2, 54, 30) |
| Right Middle Insula | BA13 |  | 9.40 | (28, 20, -4) |
| Right Anterior Insula | BA13 |  | 8.89 | (36, 16, -10) |
| Right Superior Frontal Gyrus | BA8 |  | 8.39 | (20, 32, 48) |
| Right Superior Frontal Gyrus | BA10 |  | 7.44 | (18, 66, 12) |
| Right Thalamus | Ventral Anterior |  | 6.05 | (4, -6, 4) |
| Right Nucleus Accumbens |  |  | 6.04 | (12, 6, -8) |
| Right Middle Frontal Gyrus | BA47 |  | 5.63 | (36, 36, -4) |
| **2. Left Precuneus** | **BA7** | **55080** | **-9.18** | **(-22, -62, 40)** |
| Left Inferior Parietal Lobule | BA40 |  | -6.52 | (-42, -48, 38) |
| Left Superior Parietal Lobule | BA7 |  | -6.23 | (0, -70, 60) |
| Left Superior Occipital Gyrus | BA19 |  | -5.65 | (-30, -72, 24) |
| Left Paracentral Lobule | BA4 |  | -4.74 | (0, -30, 74) |
| Right Superior Parietal Lobule | BA7 |  | -7.94 | (30, -52, 40) |
| Right Precuneus | BA7 |  | -7.08 | (14, -72, 54) |
| Right Medial Frontal Gyrus | BA6 |  | -5.81 | (2, -8, 70) |
| Right Postcentral Gyrus | BA3 |  | -5.60 | (2, -52, 68) |
| Right Superior Occipital Gyrus | BA19 |  | -5.00 | (36, -84, 22) |
| Right Postcentral Gyrus | BA40 |  | -4.92 | (64, -24, 22) |
| **3. Left Cerebellar Tuber** |  | **40552** | **-7.64** | **(-38, -66, -28)** |
| Left Cerebellar Uvula |  |  | -7.42 | (-34, -70, -26) |
| Left Fusiform Gyrus | BA37 |  | -7.32 | (-54, -58, -16) |
| Left Inferior Temporal Gyrus | BA37 |  | -6.84 | (-56, -58, -8) |
| Left Fusiform Gyrus | BA19 |  | -6.62 | (-52, -66, -12) |
| Left Cerebellar Lobule VIII |  |  | -6.06 | (-22, -62, -52) |
| Left Cerebellar Declive |  |  | -5.77 | (-14, -74, -22) |
| Left Cuneus | BA18 |  | -5.59 | (-2, -98, 4) |
| Left Fusiform Gyrus | BA18 |  | -5.01 | (-22, -92, -10) |
| Right Cerebellar Declive |  |  | -7.23 | (4, -76, -16) |
| Right Inferior Temporal Gyrus | BA37 |  | -6.63 | (56, -58, -10) |

\* = includes seed region.

| Cluster Number and Brain Region | Brodmann Area | Volume | Maximum Intensity | TLRC Coordinates |
| --- | --- | --- | --- | --- |
| Right Cerebellar Pyramis |  |  | -6.48 | (10, -72, -26) |
| Right Cerebellar Tuber |  |  | -6.19 | (46, -60, -28) |
| Right Cerebellar Uvula |  |  | -6.05 | (26, -68, -24) |
| Right Cerebellar Inferior Semi-Lunar Lobule |  |  | -5.72 | (4, -74, -46) |
| Right Fusiform Gyrus | BA37 |  | -5.66 | (44, -38, -16) |
| Right Cerebellar Lobule VIII |  |  | -5.61 | (18, -68, -50) |
| **4. Left Posterior Cingulate Cortex** | **BA31** | **33672** | **10.86** | **(-8, -58, 20)** |
| Left Precuneus | BA31 |  | 9.50 | (-8, -50, 32) |
| Left Posterior Middle Cingulate Cortex | BA24 |  | 8.90 | (-2, -22, 38) |
| Left Parahippocampal Gyrus | BA36 |  | 6.68 | (-28, -34, -8) |
| Left Hippocampus |  |  | 6.42 | (-26, -20, -14) |
| **5. Left Inferior Frontal Gyrus** | **BA9** | **11064** | **-6.56** | **(-42, 10, 30)** |
| Left Middle Frontal Gyrus | BA46 |  | -6.51 | (-42, 32, 24) |
| **6. Right Middle Frontal Gyrus** | **BA9** | **10280** | **-6.99** | **(50, 12, 32)** |
| Right Middle Frontal Gyrus | BA46 |  | -6.51 | (50, 38, 24) |
| **7. Right Posterior Insula** | **BA13** | **6832** | **6.78** | **(34, -22, 18)** |
| Right Superior Temporal Gyrus | BA41 |  | 6.28 | (50, -24, 4) |
| Right Middle Temporal Gyrus | BA21 |  | 5.60 | (58, 2, -16) |
| Right Inferior Temporal Gyrus | BA20 |  | 4.98 | (58, -14, -22) |
| **8. Right Angular Gyrus** | **BA39** | **4760** | **7.87** | **(46, -72, 36)** |
| **9. Left Angular Gyrus** | **BA39** | **3448** | **7.27** | **(-46, -74, 34)** |
| **10. Left Superior Temporal Gyrus** | **BA41** | **2960** | **5.43** | **(-54, -24, 6)** |
| **11. Left Middle Temporal Gyrus** | **BA21** | **1896** | **5.87** | **(-62, -2, -14)** |
| **12. Right Parahippocampal Gyrus** | **BA27** | **1352** | **6.74** | **(26, -32, -4)** |
| Right Hippocampus |  |  | 5.03 | (26, -16, -12) |
| **13. Right Superior Temporal Gyrus** | **BA38** | **1040** | **4.97** | **(44, 22, -26)** |
| **14. Right Superior Frontal Gyrus** | **BA6** | **920** | **-5.40** | **(30, -6, 66)** |
| **15. Right Red Nucleus** |  | **872** | **5.89** | **(2, -18, -8)** |
| **16. Left Postcentral Gyrus** | **BA3** | **632** | **4.39** | **(-38, -26, 54)** |
| **17. Right Cerebellar Lobule VIII** |  | **552** | **4.96** | **(6, -46, -50)** |
| **18. Right Middle Frontal Gyrus** | **BA10** | **416** | **-4.70** | **(46, 54, -4)** |
| **19. Right Middle Frontal Gyrus** | **BA11** | **384** | **-4.81** | **(24, 38, -10)** |
| **20. Left Middle Frontal Gyrus** | **BA10** | **288** | **-5.57** | **(-44, 52, -6)** |
| **21. Left Caudate** |  | **272** | **-4.92** | **(-18, -22, 20)** |
| **22. Right Precentral Gyrus** | **BA4** | **272** | **-4.43** | **(20, -28, 74)** |
| **23. Right Superior Frontal Gyrus** | **BA6** | **264** | **-4.05** | **(4, 4, 54)** |
| **24. Left Cerebellar Tonsil** |  | **216** | **4.56** | **(-12, -46, -40)** |

**Supplemental Table 5.** Seed: Bilateral Pregenual ACC State: Pain

| Cluster Number and Brain Region | Brodmann Area | Volume | Maximum Intensity | TLRC Coordinates |
| --- | --- | --- | --- | --- |
| **1. Left Pregenual Anterior Cingulate Cortex** | **BA32** | **116632** | **34.24** | **(-4, 40, 6)** |
| Left Caudate |  |  | 11.24 | (-14, 18, 6) |
| Left Middle Cingulate Cortex | BA24 |  | 8.90 | (-2, -22, 38) |
| Left Putamen |  |  | 8.88 | (-20, 14, -4) |
| Left Thalamus | Mediodorsal |  | 6.80 | (0, -14, 8) |
| Left Superior Frontal Gyrus | BA10 |  | 6.43 | (-24, 52, 8) |
| Left Parahippocampal Gyrus | BA28 |  | 5.78 | (-20, -14, -16) |
| Left Hippocampus |  |  | 5.49 | (-28, -32, -6) |
| Left Thalamus | Ventral posterior |  | 4.85 | (-16, -18, 2) |
| Left Thalamus | Lateroposterior |  | 4.71 | (-18, -20, 10) |
| Right Pregenual Anterior Cingulate Cortex | BA32 |  | 34.17 | (6, 42, 8) |
| Right Caudate |  |  | 12.91 | (14, 20, 4) |
| Right Superior Frontal Gyrus | BA9 |  | 9.96 | (2, 54, 30) |
| Right Anterior Insula | BA13 |  | 9.40 | (28, 20, -4) |
| Right Superior Frontal Gyrus | BA10 |  | 7.44 | (18, 66, 12) |
| Right Superior Frontal Gyrus | BA8 |  | 6.86 | (22, 38, 46) |
| Right Interpenduncular nucleus |  |  | 6.83 | (2, -18, -8) |
| Right Nucleus Accumbens |  |  | 6.70 | (12, 4, -8) |
| Right Thalamus | Mediodorsal |  | 5.86 | (4, -22, 2) |
| Right Putamen |  |  | 5.51 | (26, -2, 4) |
| Right Middle Frontal Gyrus | BA10 |  | 5.50 | (32, 48, 10) |
| Right Inferior Frontal Gyrus | BA47 |  | 5.40 | (38, 36, -6) |
| Right Claustrum |  |  | 5.36 | (32, 6, -2) |
| Right Superior Frontal Gyrus | BA6 |  | 5.28 | (12, 22, 62) |
| **2. Left Cerebellar Uvula** |  | **36808** | **-7.42** | **(-34, -70, -26)** |
| Left Fusiform Gyrus | BA37 |  | -7.32 | (-54, -58, -16) |
| Left Inferior Temporal Gyrus | BA37 |  | -6.84 | (-56, -58, -8) |
| Left Fusiform Gyrus | BA19 |  | -6.62 | (-52, -66, -12) |
| Left Cerebellar Uvula |  |  | -6.23 | (-22, -66, -24) |
| Left Cerebellar Declive |  |  | -5.88 | (-2, -80, -18) |
| Left Cerebellar Culmen |  |  | -5.36 | (-28, -54, -26) |
| Left Cuneus | BA18 |  | -5.31 | (-4, -90, 16) |
| Left Lingual Gyrus | BA18 |  | -5.08 | (-2, -94, -12) |
| Left Middle Occipital Gyrus | BA18 |  | -4.77 | (-34, -88, 2) |
| Right Fusiform Gyrus | BA19 |  | -6.45 | (40, -72, -18) |
| Right Cerebellar Declive |  |  | -6.44 | (14, -72, -20) |
| Right Middle Temporal Gyrus | BA37 |  | -6.30 | (56, -54, -10) |

\* = includes seed region.

| Cluster Number and Brain Region | Brodmann Area | Volume | Maximum Intensity | TLRC Coordinates |
| --- | --- | --- | --- | --- |
| Right Fusiform Gyrus | BA37 |  | -6.00 | (44, -56, -18) |
| Right Cerebellar Lobule VIII |  |  | -5.24 | (14, -70, -52) |
| Right Cerebellar Culmen |  |  | -4.71 | (6, -62, -6) |
| Right Cerebellar Uvula |  |  | -4.52 | (32, -62, -26) |
| Right Cerebellar Fastigium |  |  | -4.35 | (8, -56, -20) |
| **3. Left Posterior Cingulate Cortex** | **BA31** | **29800** | **9.71** | **(-8, -60, 20)** |
| Left Precuneus | BA31 |  | 9.50 | (-8, -50, 32) |
| Left Posterior Cingulate Cortex | BA30 |  | 8.71 | (-8, -56, 10) |
| **4. Right Superior Parietal Lobule** | **BA7** | **28208** | **-7.29** | **(30, -52, 40)** |
| Right Precuneus | BA7 |  | -6.07 | (26, -66, 32) |
| Left Inferior Parietal Lobule | BA40 |  | -6.52 | (-42, -48, 38) |
| Left Precuneus | BA7 |  | -6.05 | (-26, -64, 32) |
| Left Angular Gyrus | BA7 |  | -5.54 | (-30, -56, 40) |
| Left Supramarginal Gyrus | BA40 |  | -4.95 | (-54, -42, 32) |
| **5. Left Middle Frontal Gyrus** | **BA9** | **9752** | **-6.04** | **(-50, 12, 32)** |
| Left Middle Frontal Gyrus | BA6 |  | -5.59 | (-50, 6, 44) |
| **6. Left Middle Frontal Gyrus** | **BA8** | **6720** | **10.05** | **(-22, 36, 42)** |
| **7. Right Middle Frontal Gyrus** | **BA9** | **4320** | **-4.72** | **(50, 12, 30)** |
| Right Middle Frontal Gyrus | BA8 |  | -4.67 | (52, 10, 38) |
| Right Superior Frontal Gyrus | BA9 |  | -4.51 | (40, 36, 32) |
| **8. Right Inferior Parietal Lobule** | **BA40** | **4072** | **-5.86** | **(56, -40, 28)** |
| **9. Right Angular Gyrus** | **BA39** | **2904** | **7.06** | **(50, -70, 30)** |
| Right Middle Temporal Gyrus | BA37 |  | 4.78 | (54, -62, 18) |
| **10. Right Parahippocampal Gyrus** | **BA27** | **1632** | **6.09** | **(26, -34, -4)** |
| Right Hippocampus |  |  | 5.62 | (28, -22, -14) |
| **11. Right Posterior Insula** | **BA13** | **1568** | **5.80** | **(34, -22, 14)** |
| Right Superior Temporal Gyrus | BA41 |  | 5.23 | (48, -22, 8) |
| **12. Left Posterior Insula** | **BA13** | **1464** | **4.79** | **(-48, -14, 12)** |
| **13. Left Angular Gyrus** | **BA39** | **1464** | **6.31** | **(-44, -76, 36)** |
| **14. Left Cerebellar Inferior Semilunar Lobule** |  | **1456** | **-5.34** | **(-6, -72, -46)** |
| Left Cerebellar Lobule VIII |  |  | -4.81 | (-26, -68, -52) |
| **15. Right Middle Occipital Gyrus** | **BA19** | **1104** | **-5.03** | **(28, -92, 20)** |
| **16. Left Cerebellar Tonsil** |  | **856** | **5.68** | **(-4, -48, -48)** |
| **17. Right Cerebellar Tonsil** |  | **616** | **4.93** | **(12, -46, -40)** |
| **18. Right Superior Frontal Gyrus** | **BA6** | **472** | **-4.56** | **(30, -4, 66)** |
| **19. Right Inferior Frontal Gyrus** | **BA10** | **352** | **-5.60** | **(46, 54, 0)** |
| **20. Left Superior Parietal Lobule** | **BA7** | **312** | **-4.86** | **(-24, -54, 66)** |
| **21. Nucleus Prepositus** |  | **208** | **-5.13** | **(0, -36, -34)** |
| **22. Left Cerebellar Declive** |  | **200** | **-4.41** | **(-8, -64, -20)** |
| **23. Left Pontine Nuclei** |  | **80** | **3.74** | **(-4, -24, -40)** |

**Supplemental Table 6.** Seed: Bilateral Subgenual ACC State: Control

| Cluster Number and Brain Region | Brodmann Area | Volume | Maximum Intensity | TLRC Coordinates |
| --- | --- | --- | --- | --- |
| **1. Right Subgenual Anterior Cingulate Cortex** | **BA24** | **81848** | **35.06** | **(4, 28, -4)** |
| Left Subgenual Anterior Cingulate Cortex | BA24 |  | 34.59 | (-4, 30, 0) |
| Left Medial Frontal Gyrus | BA10 |  | 12.40 | (-2, 60, 14) |
| Left Superior Frontal Gyrus | BA8 |  | 8.51 | (-22, 28, 48) |
| Left Middle Frontal Gyrus | BA10 |  | 5.16 | (-22, 60, 22) |
| Right Superior Frontal Gyrus | BA6 |  | 9.12 | (20, 34, 56) |
| Right Anterior Insula | BA13 |  | 6.98 | (28, 20, -10) |
| Right Inferior Frontal Gyrus | BA47 |  | 6.52 | (20, 34, -6) |
| **2. Left Precuneus** | **BA31** | **25064** | **9.72** | **(-8, -52, 32)** |
| Left Posterior Cingulate Cortex | BA31 |  | 9.28 | (-8, -44, 36) |
| Left Parahippocampal Gyrus | BA34 |  | 7.26 | (-20, -12, -16) |
| Left Parahippocampal Gyrus | BA30 |  | 7.12 | (-10, -48, 6) |
| Right Posterior Cingulate Cortex | BA31 |  | 7.45 | (4, -56, 24) |
| **3. Left Cerebellar Culmen** |  | **13216** | **-6.81** | **(-34, -60, -26)** |
| Left Cerebellar Declive |  |  | -6.61 | (-8, -72, -20) |
| Left Cerebellar Pyramis |  |  | -5.97 | (-8, -78, -24) |
| Left Cerebellar Tonsil |  |  | -5.75 | (-36, -46, -34) |
| Left Cerebellar Tuber |  |  | -5.67 | (-42, -54, -24) |
| Right Cerebellar Uvula |  |  | -5.74 | (28, -68, -26) |
| Right Lingual Gyrus |  |  | -4.32 | (2, -88, -18) |
| **4. Right Precuneus** | **BA7** | **9328** | **-5.86** | **(10, -74, 34)** |
| Right Superior Parietal Lobule | BA7 |  | -4.73 | (14, -62, 62) |
| Left Precuneus | BA7 |  | -5.69 | (-8, -72, 52) |
| Left Superior Parietal Lobule | BA7 |  | -5.31 | (-12, -68, 54) |
| Left Cuneus | BA19 |  | -5.12 | (-14, -82, 42) |
| **5. Right Middle Temporal Gyrus** | **BA21** | **8224** | **7.31** | **(58, -8, -12)** |
| Right Superior Temporal Gyrus | BA38 |  | 6.36 | (40, 20, -28) |
| Right Superior Temporal Gyrus | BA21 |  | 6.27 | (54, 6, -14) |
| **6. Right Inferior Frontal Gyrus** | **BA9** | **8176** | **-6.20** | **(42, 6, 22)** |
| Right Inferior Frontal Gyrus | BA44 |  | -5.87 | (50, 8, 18) |
| Right Anterior Insula | BA13 |  | -5.80 | (34, 16, 18) |
| **7. Right Middle Frontal Gyrus** | **BA10** | **7792** | **-6.54** | **(36, 38, 22)** |
| Right Middle Frontal Gyrus | BA9 |  | -6.26 | (36, 48, 36) |
| **8. Right Superior Frontal Gyrus** | **BA6** | **6800** | **-6.50** | **(4, 6, 52)** |
| Left Superior Frontal Gyrus | BA6 |  | -4.77 | (-16, -2, 70) |
| **9. Right Supramarginal Gyrus** | **BA40** | **6520** | **-6.61** | **(62, -42, 32)** |
| Right Inferior Parietal Lobule | BA40 |  | -5.95 | (56, -40, 42) |

\* = includes seed region.

| Cluster Number and Brain Region | Brodmann Area | Volume | Maximum Intensity | TLRC Coordinates |
| --- | --- | --- | --- | --- |
| Right Superior Parietal Lobule | BA7 |  | -4.76 | (38, -56, 54) |
| **10. Right Angular Gyrus** | **BA39** | **6144** | **9.28** | **(46, -70, 34)** |
| **11. Left Middle Temporal Gyrus** | **BA21** | **5704** | **7.60** | **(-54, -4, -12)** |
| Left Superior Temporal Gyrus | BA38 |  | 5.03 | (-46, 14, -28) |
| **12. Left Angular Gyrus** | **BA39** | **5192** | **8.34** | **(-46, -72, 36)** |
| **13. Left Inferior Parietal Lobule** | **BA40** | **5048** | **-5.51** | **(-62, -36, 34)** |
| Left Superior Parietal Lobule | BA7 |  | -5.07 | (-38, -58, 48) |
| **14. Right Hippocampus** |  | **4856** | **7.04** | **(26, -16, -14)** |
| Right Parahippocampal Gyrus | BA36 |  | 6.90 | (24, -32, -10) |
| **15. Left Middle Insula** | **BA13** | **4488** | **-5.87** | **(-46, 4, 14)** |
| Left Inferior Frontal Gyrus | BA44 |  | -5.34 | (-42, 16, 14) |
| **16. Left Postcentral Gyrus** | **BA2** | **4208** | **5.44** | **(-44, -26, 42)** |
| Left Precentral Gyrus | BA6 |  | 5.42 | (-56, -12, 38) |
| **17. Left Parahippocampal Gyrus** | **BA28** | **4160** | **7.10** | **(-22, -18, -14)** |
| Left Hippocampus |  |  | 6.30 | (-24, -14, -16) |
| **18. Left Inferior Frontal Gyrus** | **BA47** | **4056** | **7.86** | **(-22, 30, -10)** |
| **19. Left Superior Frontal Gyrus** | **BA9** | **3760** | **-6.45** | **(-38, 44, 30)** |
| Left Middle Frontal Gyrus | BA10 |  | -4.15 | (-40, 42, 12) |
| **20. Right Thalamus** | **Ventrolateral** | **1184** | **-5.02** | **(16, -12, 16)** |
| Right Thalamus | Lateroposterior |  | -4.77 | (20, -18, 10) |
| **21. Right Precentral Gyrus** | **BA4** | **544** | **4.71** | **(30, -30, 50)** |
| **22. Right Cuneus** | **BA18** | **496** | **-4.44** | **(26, -74, 24)** |
| **23. Right Cerebellar Semi-Lunar Lobule** |  | **376** | **4.19** | **(24, -82, -38)** |
| **24. Right Rostral Ventral Medulla** |  | **368** | **-4.52** | **(4, -22, -50)** |
| **25. Right Postcentral Gyrus** | **BA2** | **264** | **4.04** | **(48, -24, 50)** |
| **26. Left Cerebellar Lobule VIII** |  | **232** | **-4.47** | **(-22, -30, -48)** |
| **27. Right Thalamus** | **Pulvinar** | **120** | **-5.47** | **(18, -32, 14)** |
| **28. Left Interpenduncular Nucleus** |  | **88** | **4.84** | **(-2, -20, -16)** |

**Supplemental Table 7.** Seed: Bilateral Subgenual ACC State: Pain

| Cluster Number and Brain Region | Brodmann Area | Volume | Maximum Intensity | TLRC Coordinates |
| --- | --- | --- | --- | --- |
| **1. Left Subgenual Anterior Cingulate Cortex** | **BA24** | **85888** | **34.59** | **(-4, 30, 0)** |
| Left Medial Frontal Gyrus | BA10 |  | 11.76 | (-2, 60, 14) |
| Left Superior Frontal Gyrus | BA8 |  | 7.86 | (-18, 32, 46) |
| Left Superior Frontal Gyrus | BA10 |  | 7.32 | (-4, 66, 30) |
| Left Parahippocampal Gyrus | BA34 |  | 7.26 | (-20, -12, -16) |
| Left Parahippocampal Gyrus | BA36 |  | 7.24 | (-26, -32, -16) |
| Left Hippocampus |  |  | 7.07 | (-24, -14, -16) |
| Left Middle Frontal Gyrus | BA6 |  | 5.03 | (-30, 16, 46) |
| Right Superior Frontal Gyrus | BA8 |  | 8.37 | (20, 32, 46) |
| Right Superior Frontal Gyrus | BA6 |  | 8.04 | (20, 34, 56) |
| Right Hippocampus |  |  | 7.31 | (26, -16, -14) |
| Right Superior Frontal Gyrus | BA10 |  | 6.73 | (16, 60, 18) |
| Right Inferior Frontal Gyrus | BA47 |  | 6.25 | (30, 24, -8) |
| Right Anterior Insula | BA13 |  | 6.10 | (42, 38, -10) |
| Right Middle Frontal Gyrus | BA11 |  | 6.01 | (24, 38, -2) |
| Right Lateral Globus Pallidus |  |  | 5.99 | (20, -4, -8) |
| Right Middle Insula | BA13 |  | 5.92 | (30, 12, -8) |
| Right Medial Globus Pallidus |  |  | 5.72 | (10, 0, -8) |
| **2. Left Posterior Cingulate Cortex** | **BA31** | **25168** | **9.28** | **(-8, -44, 36)** |
| Left Posterior Cingulate Cortex | BA30 |  | 6.94 | (-8, -54, 10) |
| Right Posterior Cingulate Cortex | BA31 |  | 8.77 | (4, -56, 24) |
| **3. Right Middle Temporal Gyrus** | **BA21** | **7408** | **6.80** | **(58, -4, -12)** |
| Right Superior Temporal Gyrus | BA38 |  | 6.56 | (42, 26, -26) |
| **4. Right Middle Frontal Gyrus** | **BA10** | **7024** | **-6.22** | **(38, 38, 18)** |
| Right Middle Frontal Gyrus | BA9 |  | -5.74 | (36, 48, 36) |
| **5. Right Supramarginal Gyrus** | **BA40** | **6992** | **-6.25** | **(58, -40, 32)** |
| Right Superior Parietal Lobule | BA7 |  | -5.87 | (40, -54, 54) |
| Right Inferior Parietal Lobule | BA40 |  | -5.30 | (32, -48, 38) |
| Right Cuneus | BA19 |  | -4.70 | (30, -70, 28) |
| **6. Left Middle Temporal Gyrus** | **BA21** | **5360** | **6.75** | **(-54, -6, -12)** |
| **7. Left Angular Gyrus** | **BA39** | **4912** | **7.15** | **(-42, -76, 32)** |
| **8. Right Inferior Frontal Gyrus** | **BA9** | **4024** | **-5.89** | **(46, 10, 24)** |
| **9. Right Angular Gyrus** | **BA39** | **3904** | **8.17** | **(48, -70, 26)** |
| **10. Left Inferior Frontal Gyrus** | **BA47** | **3888** | **6.03** | **(-22, 30, -10)** |
| Left Middle Frontal Gyrus | BA11 |  | 5.67 | (-40, 34, -10) |
| **11. Right Cerebellar Pyramis** |  | **3456** | **-5.59** | **(36, -66, -32)** |
| Right Cerebellar Culmen |  |  | -4.44 | (38, -50, -30) |

\* = includes seed region.

| **Cluster Number and Brain Region** | **Brodmann Area** | **Volume** | **Maximum Intensity** | **TLRC Coordinates** |
| --- | --- | --- | --- | --- |
| **12. Left Superior Frontal Gyrus** | **BA9** | **2760** | **-5.51** | **(-42, 40, 32)** |
| **13. Left Cerebellar Uvula** |  | **2592** | **-5.73** | **(-8, -70, -32)** |
| Right Cerebellar Uvula |  |  | -4.49 | (6, -70, -32) |
| **14. Left Inferior Frontal Gyrus** | **BA44** | **1816** | **-5.11** | **(-52, 12, 14)** |
| **15. Right Anterior Insula** | **BA13** | **1656** | **-4.96** | **(32, 16, 12)** |
| **16. Left Cerebellar Tonsil** |  | **1464** | **-4.61** | **(-22, -64, -32)** |
| **17. Left Anterior Insula** | **BA13** | **1344** | **-5.06** | **(-36, 12, 16)** |
| **18. Left Middle Occipital Gyrus** | **BA19** | **680** | **-4.78** | **(-30, -76, 20)** |
| **19. Left Superior Parietal Lobule** | **BA7** | **352** | **-4.08** | **(-20, -70, 54)** |
| **20. Left Thalamus** | **Lateroposterior** | **336** | **-4.64** | **(-20, -18, 16)** |
| **21. Right Lateral Globus Pallidus** |  | **264** | **-4.24** | **(24, -16, 6)** |
| **22. Left Cerebellar Inferior Semi-Lunar Lobule** |  | **216** | **4.29** | **(-12, -78, -40)** |
| **23. Right Cerebellar Declive** |  | **144** | **-4.28** | **(4, -70, -18)** |
| **24. Right Caudate** |  | **144** | **-4.50** | **(18, -8, 16)** |

**Supplemental Table 8.** Seed: Periaqueductal Gray State: Control

| Cluster Number and Brain Region | Brodmann Area | Volume | Maximum Intensity | TLRC Coordinates |
| --- | --- | --- | --- | --- |
| **1. Periaqueductal Gray\*** |  | **19568** | **19.52** | **(-2, -32, -10)** |
| Left Cerebellar Culmen |  |  | 8.46 | (-6, -52, -8) |
| Right Thalamus | Anterior Nucleus |  | 6.93 | (2, 4, 6) |
| Left Thalamus | Anterior Nucleus |  | 6.86 | (-6, -4, 2) |
| Right Cerebellar Culmen |  |  | 8.59 | (4, -50, -12) |
| Right Subthalamic Nucleus |  |  | 6.62 | (16, -14, -4) |
| Right Putamen |  |  | 5.60 | (28, -12, -4) |
| Right Cerebellar Declive |  |  | 4.71 | (8, -62, -18) |
| **2. Right Middle Frontal Gyrus** | **BA47** | **816** | **-5.64** | **(46, 44, -6)** |
| **3. Left Pregenuel Anterior Cingulate Cortex** | **BA24** | **608** | **5.82** | **(-2, 34, 12)** |
| **4. Left Ventromedial Prefrontal Gyrus** | **BA10** | **384** | **-5.41** | **(-2, 44, -10)** |
| **5. Right Middle Insula** | **BA13** | **320** | **4.38** | **(42, 2, 10)** |
| **6. Left Parahippocampal Gyrus** | **BA28** | **296** | **5.21** | **(-16, -28, -14)** |
| **7. Right Middle Frontal Gyrus** | **BA11** | **280** | **-4.66** | **(44, 46, -10)** |
| **8. Left Middle Frontal Gyrus** | **BA10** | **272** | **-4.52** | **(-28, 48, -6)** |
| **9. Right Cerebellar Tonsil** |  | **192** | **4.80** | **(36, -40, -34)** |
| **10. Left Nucleus Raphe Magnus** |  | **88** | **4.44** | **(-4, -28, -54)** |
| **11. Right Dorsolateral Pons** |  | **64** | **4.18** | **(10, -22, -20)** |

\* = includes seed region.

**Supplemental Table 9.** Seed: Periaqueductal Gray State: Pain

| Cluster Number and Brain Region | Brodmann Area | Volume | Maximum Intensity | TLRC Coordinates |
| --- | --- | --- | --- | --- |
| **1. Periaqueductal Gray\*** |  | **38536** | **19.08** | **(2, -32, -10)** |
| Left Subthalamic Nucleus |  |  | 7.40 | (-10, -14, -2) |
| Left Thalamus | Anterior Nucleus |  | 6.86 | (-6, -4, 2) |
| Left Anterior Insula | BA13 |  | 5.73 | (-38, 14, -2) |
| Left Caudate |  |  | 5.48 | (-12, 8, 6) |
| Left Lateral Globus Pallidus |  |  | 5.34 | (-16, 6, 2) |
| Left Putamen |  |  | 5.10 | (-26, 12, 12) |
| Right Subthalamic Nucleus |  |  | 6.62 | (16, -14, -4) |
| Right Cerebellar Culmen |  |  | 6.27 | (12, -44, -20) |
| Right Thalamus | Ventrolateral Nucleus |  | 6.09 | (10, -8, 8) |
| Right Thalamus | Mediodorsal Nucleus |  | 6.04 | (8, -16, 14) |
| Right Putamen |  |  | 5.82 | (24, 8, 4) |
| Right Caudate |  |  | 5.25 | (10, 2, 10) |
| Right Cerebellar Tonsil |  |  | 5.21 | (2, -48, -34) |
| Right Lateral Globus Pallidus |  |  | 4.93 | (26, -20, -2) |
| Right Inferior Frontal Gyrus | BA45 |  | 4.50 | (42, 22, 12) |
| **2. Left Pregenuel Anterior Cingulate Cortex** | **BA24** | **1080** | **5.34** | **(-8, 38, 4)** |
| Right Pregenuel Anterior Cingulate Cortex | BA24 |  | 5.25 | (2, 34, 12) |
| **3. Right Inferior Occipital Gyrus** | **BA18** | **856** | **-5.42** | **(36, -90, -10)** |
| **4. Left Ventromedial Frontal Gyrus** | **BA10** | **680** | **-4.80** | **(-2, 46, -10)** |
| **5. Left Superior Frontal Gyrus** | **BA9** | **648** | **-5.30** | **(-32, 52, 26)** |
| **6. Left Cerebellar Uvula** |  | **536** | **4.73** | **(0, -68, -30)** |
| **7. Right Precentral Gyrus** | **BA6** | **440** | **-4.72** | **(30, -14, 68)** |
| **8. Right Cerebellar Tonsil** |  | **424** | **4.90** | **(12, -44, -48)** |
| **9. Left Parahippocampal Gyrus** | **BA36** | **368** | **4.66** | **(-26, -32, -10)** |
| **10. Left Middle Temporal Gyrus** | **BA22** | **360** | **-5.00** | **(-58, -32, 2)** |
| **11. Right Inferior Parietal Lobule** | **BA40** | **312** | **-4.32** | **(48, -30, 50)** |
| **12. Left Cuneus** | **BA19** | **280** | **-4.53** | **(-24, -82, 30)** |
| **13. Right Inferior Parietal Lobule** | **BA40** | **272** | **-5.13** | **(38, -38, 38)** |
| **14. Left Cerebellar Tonsil** |  | **216** | **4.56** | **(-16, -32, -44)** |
| **15. Right Subgenual Anterior Cingulate Cortex** | **BA25** | **144** | **-4.28** | **(6, 14, -8)** |
| **16. Left Putamen** |  | **144** | **3.88** | **(-26, -6, -6)** |
| **17. Right Rostral Ventral Medulla** |  | **80** | **4.34** | **(6, -28, -48)** |

\* = includes seed region.

